# A novel post-translational modification of fimbriae drives pathogenicity in *Klebsiella pneumoniae*

**DOI:** 10.1101/2025.10.03.680091

**Authors:** Genevieve Dobihal, Kristen Lewis, Lisa Yu, Carmen Herrera, M Stephen Trent, Josué Flores Kim, Anne-Catrin Uhlemann

## Abstract

Multi-drug resistant Gram-negative bacteria, including carbapenem-resistant *Klebsiella pneumoniae* (CR*Kp*), are a public health emergency. The predominant CR*Kp* sequence type worldwide is ST258. However, the factors underlying ST258’s epidemic success are not well defined. The understudied two-component system CrrAB is a genomic feature of ST258 and has been hypothesized to contribute to its global dominance. Despite this, the molecular details underpinning CrrAB’s contribution to ST258 pathogenicity are not well understood. We used RNA-sequencing to identify the regulon of CrrA and found that CrrAB induces the expression of a novel gene, encoding Crr-regulated fimbriae modifying protein (CfmP). CfmP post-translationally modifies fimbriae to significantly increase host cell adhesion and high bacterial loads within the host, consequently increasing ST258 virulence. CrrAB also drives high antibiotic resistance in CR*Kp*. Thus, our data places CrrAB at the intersection of virulence and antibiotic resistance supporting its function as an important regulatory system driving the pathogenicity of ST258.

---

Multi-drug resistant (MDR) Gram-negative bacterial infections are among the most urgent health threats^1^. *Klebsiella pneumoniae* (*Kp*) is a major cause of MDR infections, accounting for the third most global deaths attributable to or associated with bacterial MDR^1^. Especially concerning is carbapenem resistant *Kp* (CR*Kp*), which is a leading cause of invasive, drug-resistant infections in many human body sites, with high rates of morbidity and mortality^2–8^. Resistance has emerged in many *Kp* clonal groups, but only a few lineages have come to dominate MDR *Kp* infections worldwide. In particular, the global spread of CR*Kp* has been driven in main part by sequence type 258 (ST258)^2,8,9^. ST258 represents a particular challenge for healthcare providers, as it exhibits high levels of multi-class drug resistance, severely limiting treatment options^8^. Notably, the reasons underlying ST258’s world-wide dissemination and the molecular mechanisms underpinning its pathogenic success are unclear^2,8,9^.

Genomic analyses performed on ST258 isolates have identified several loci that are associated with pathogenicity in ST258^4,10–15^. However, molecular studies to determine the relevance of these potential virulence determinants are lacking. Notably, the understudied two component system (TCS) CrrAB was recently identified as a genomic feature of ST258^16,17^ and has been suggested to confer pathogenic and physiological benefits to ST258^17,18^. Further, missense mutations in CrrAB were identified in clinical isolates of ST258 CR*Kp*, which were found to confer high antibiotic resistance and high virulence^19–22^. In particular, these CrrAB mutations induce resistance to last-line polymyxin antibiotics, cationic non-ribosomal peptides that disrupt the bacterial cell envelope. CrrAB was shown to induce polymyxin resistance (PR), in part, through the activation of *eptA* and the *arn* operon. These genes encode enzymes that catalyze the addition of modifications to the lipopolysaccharide (LPS), the primary surface molecule of Gram-negative bacteria, protecting against polymyxin attack^19–22^. Intriguingly, polymyxin antibiotics function similarly to host-produced antimicrobial peptides (AMPs). This suggests that CrrAB may function in ST258’s response to the host environment. However, EptA- and Arn-driven LPS modifications do not fully correlate with increased virulence in *Kp*, suggesting that CrrAB functions to increase ST258 pathogenicity through additional, unknown mechanisms^22^.

Here, we report that a previously uncharacterized gene in the CrrA regulon, *cfmP*, drives virulence in ST258 *Kp.* We show that CfmP induces the addition of a novel post-translational modification to the major pilin FimA of type I pili, or fimbriae, on the cell surface. These modified fimbriae lead to increased host cell attachment and bacterial proliferation within the host, to cause high pathogenicity. Overall, these findings expand our mechanistic understanding of how CrrAB, a genomic feature of ST258, contributes to the pathogenicity of this high-risk clonal group.

## Results

### *crrAB* is conserved in ST258 CR*Kp*

ST258 has been the dominant driver of CR*Kp* infections globally. The CrrAB TCS (Fig. 1a) was previously reported to be a genomic feature of ST258^16,17^, suggesting that it may contribute to its pathogenic success. We analyzed the genomes of a collection of 918 CR*Kp* strains isolated at Columbia University Irving Medical Center^23–25^. We found that 58% belonged to 258 clonal group (CG258), containing ST258 and its single-locus variants ST11, ST340, and ST512, while the remaining 383 isolates (42%) were distributed amongst 125 different sequence types (Fig. 1a). Among the ST258 isolates, we performed genomic searches for *crrA* and *crrB*. Out of 249 isolates, 218 contained both genes (Fig. 1a). Nineteen genomes were lacking *crrAB* but contained genes in the immediate vicinity of *crrAB*, and had insertion sequences (IS) nearby, suggesting IS-driven transpositions may have removed *crrAB*. The genomes completely lacking *crrAB* and its nearby genes were in a small cluster of closely-related strains. Overall, 95% of the ST258 isolates we searched contained *crrAB* or genes in its surrounding locus, supporting that CrrAB is a conserved feature of ST258.

**Fig. 1:**
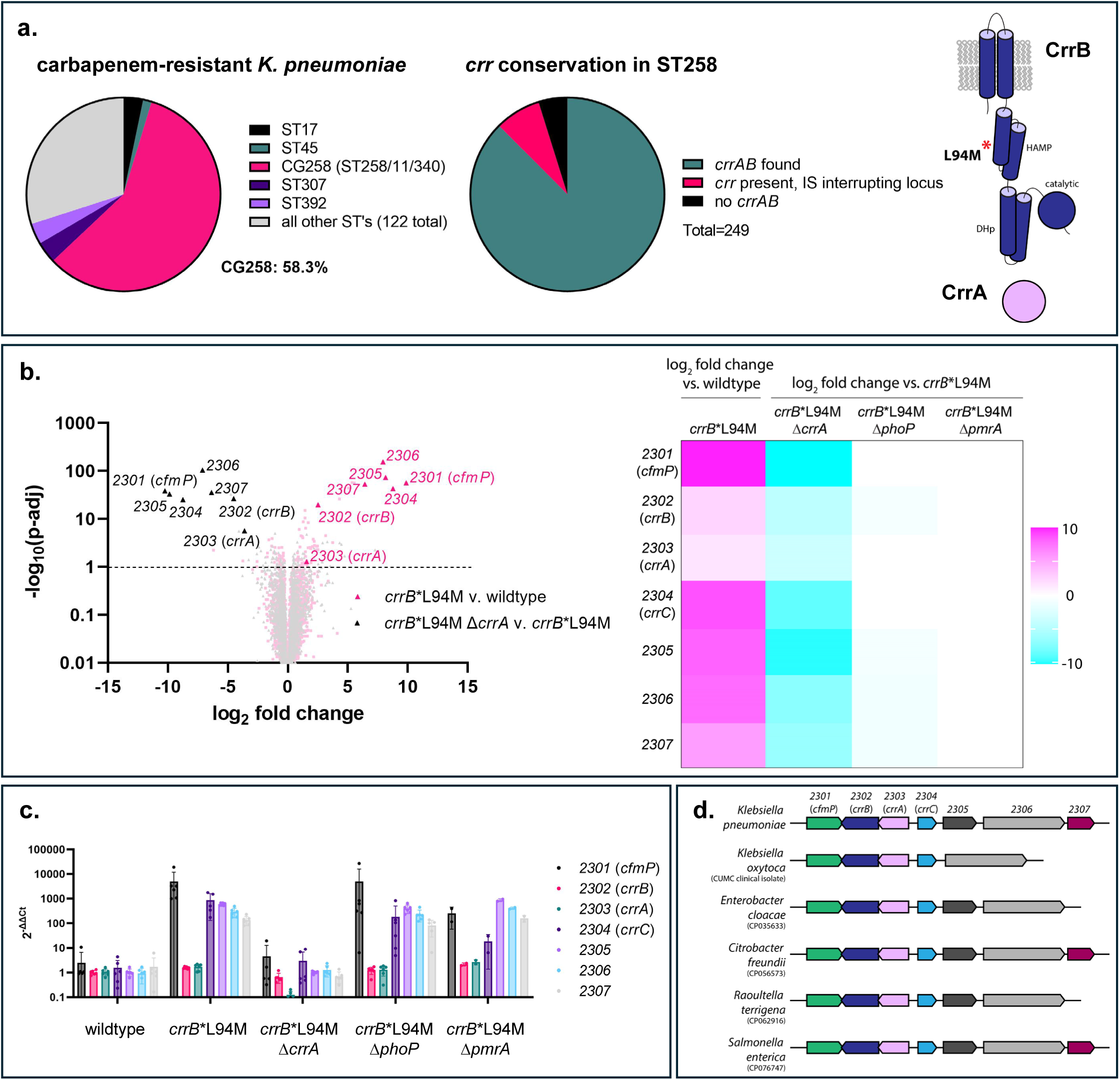
CrrAB transcriptionally activates five uncharacterized genes in its surrounding locus. **(a)** Genomic searches performed on CR*Kp* isolates collected from CUIMC (n=918). 535 isolates belonged to CG258 (ST258, ST11, ST340, ST512), while the rest (n=383) were distributed among 126 different sequence types. Among ST258 isolates collected from unique patients (n=249), a majority (n=237) contained *crrAB* or genes in the *crr* locus. **(b)** Volcano plot (left) of RNA-seq data generated from comparing the transcriptional profile of the clinical mutant *crrB*\*L94M vs. wildtype (pink) and *crrB*\*L94M *ΔcrrA* vs. *crrB*\*L94M (grey). Genes in the *crr* locus are highlighted. Heat map (right) of the log_2_ fold change of the *crr* locus in the indicated strains. **(c)** Quantitative reverse transcription PCR (RT-qPCR) was used to confirm the fold change in transcription of the genes in the *crr* locus in the indicated strains. **(d)** A schematic of the *crr* loci of *Kp* and other *Enterobacteriaceae* species. Searches were performed using publicly-available genomes (GenBank accession numbers are noted in parentheses). CrrB was used to generate the sequence similarity network used for genomic neighborhood analyses of the *crr* locus.

### CrrAB upregulates its surrounding *crr* locus

The CrrAB TCS was previously shown to induce high virulence in *Kp*^22^. CrrB is a membrane-anchored sensor histidine kinase, while CrrA is a DNA-binding response regulator. To identify the full landscape of transcriptional changes induced by CrrAB and the specific genes involved in the increased pathogenicity driven by CrrAB, we performed RNA sequencing (RNAseq). We compared the transcriptomes of wildtype *Kp* and an isogenic strain into which we introduced a clinically-identified, hyperactive mutant of CrrB, *crrB*\*L94M^22,23^ (Fig. 1a,b). It has been shown that CrrAB and two other TCSs, PhoPQ and PmrAB, share transcriptional targets in *Kp*^19,22^. Therefore, to determine CrrAB-specific regulation, we compared the transcriptional profiles of *crrB*\*L94M in the absence of the response regulators of each individual signaling system (*crrB*\*L94M Δ*crrA*, crrB*L94M Δ*pmrA*, and *crrB*\*L94M Δ*phoP*). In agreement with previous studies, our results showed that *crrB*\*L94M, in coordination with PmrAB and PhoPQ, increases transcription of the *arn* operon and *eptA* to drive PR^26^ (Extended Data Fig. 1). Intriguingly, we found that CrrA, but not PhoP or PmrA, leads to the dramatic upregulation of five genes of unknown function in the genomic locus surrounding *crrAB*, which we have termed the *crr* locus (Fig. 1b,c). These genes were previously identified to be associated with *crrAB* in *Klebsiella* species and in some *Citrobacter* and *Enterobacter* species^16,17^, although their roles in the pathogenicity of *Kp* and other *Enterobacteriaceae* are unknown. Gene neighborhood analysis was performed^27,28^ and further identified the *crr* locus in diverse *Enterobacteriaceae* species, including several human pathogens (Fig. 1d), suggesting that it may contribute to CrrAB’s function in virulence. Importantly, we found that the same CrrA regulon identified in our isogenic strains is also upregulated in an original clinical *Kp* isolate identified harboring *crrB*\*L94M^22,23^ (Extended Data Fig. 4a).

### CrrAB-regulated *cfmP* confers high virulence in *crrB*\*L94M

We next investigated whether genes in the *crr* locus are responsible for the drug resistance and virulence phenotypes observed in CrrAB clinical mutants^22^. As previously shown, CrrC is necessary for PR in *crrB*\*L94M (Fig. 2a). *crrC* is directly upstream of *crrA* and suggested to encode a connector protein, mediating signaling cross-talk between CrrAB and PmrAB^19,22^. Using the *Galleria mellonella* virulence model, we found that only one of the genes in the *crr* locus, *cfmP*, completely reduced the high virulence of *crrB*\*L94M back to wildtype levels (Fig. 2b). To further investigate the mechanisms underlying CrrAB and *cfmP*-driven virulence, we determined the bacterial loads over the course of *G. mellonella* infections. We showed that, in addition to causing high mortality of *G. mellonella* (Fig. 2b,c), CrrAB’s regulation of *cfmP* causes high bacterial proliferation in the host (Fig. 2c). Further, we found that overexpression of *cfmP* increases virulence and bacteria loads in wildtype *Kp* (Fig. 2d). Importantly, virulence was restored when *cfmP* was complemented *in trans* (Fig. 2e, Extended Data Fig. 2a), and the decrease in virulence of *crrB*\*L94M Δ*cfmP* was not due to off-target effects on the transcription of the *crr* locus (Extended Data Fig. 2b). Further, we found that deletion of *cfmP* in the original *Kp* clinical isolate harboring the *crrB*\*L94M mutation also completely decreases the virulence of this isolate (Extended Data Fig. 4b).

**Fig. 2:**
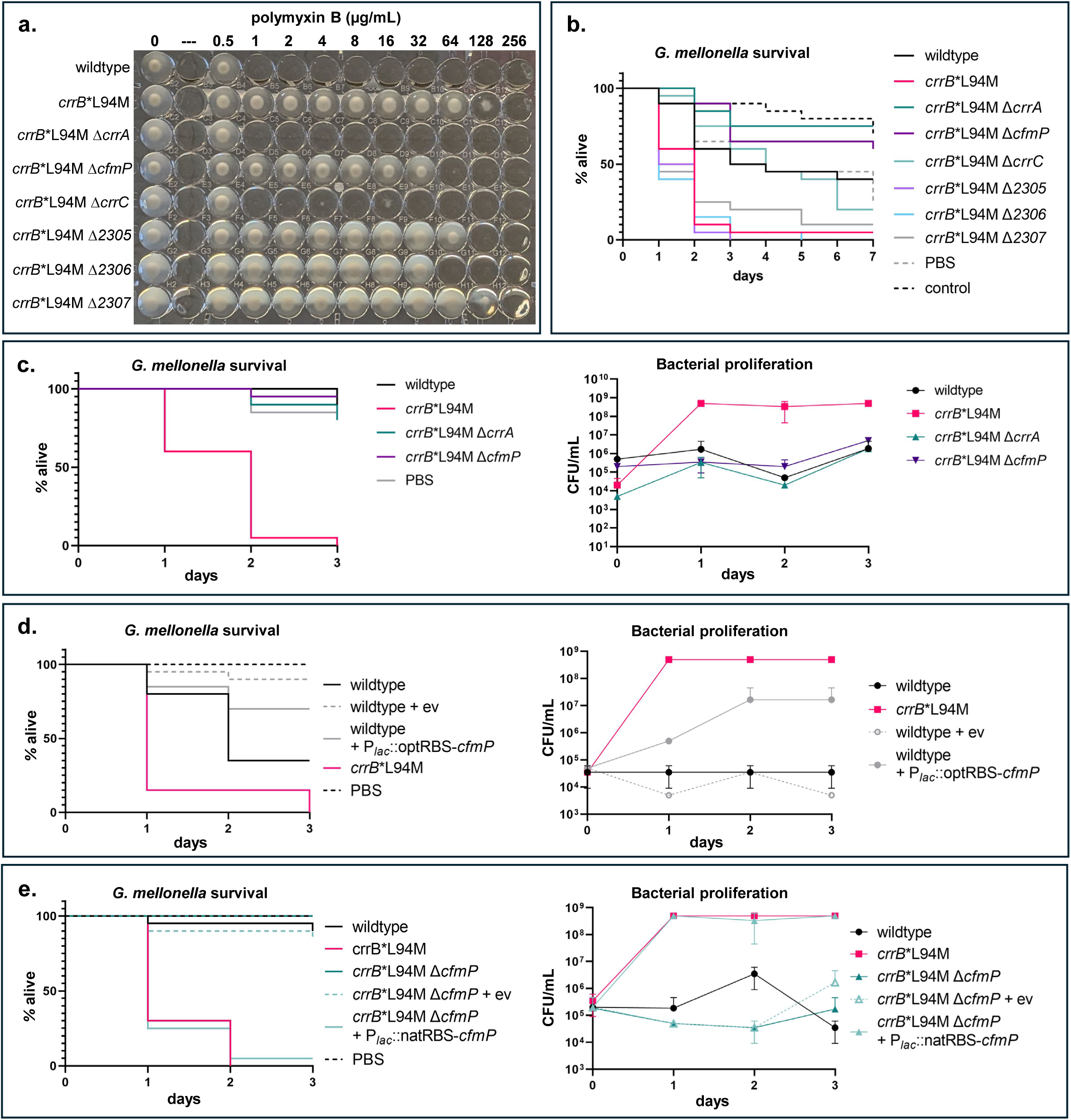
A novel gene in the CrrA regulon and *crr* locus, *cfmP*, drives the high virulence of CrrB*L94M. **(a)** A polymyxin B broth microdilution (BMD) plate of the indicated mutants. **(b)** *G. mellonella* infections using the indicated strains. **(c)** *G. mellonella* survival curve and *G. mellonella* bacterial load quantification of infections with wildtype, *crrB*\*L94M, *crrB*\*L94M Δ*crrA*, and *crrB*\*L94M Δ*cfmP.* **(d)** *G. mellonella* infections of wildtype, *crrB*\*L94M, and wildtype harboring empty vector (ev) or overexpression plasmids (pUC19-P*_lac_*::optRBS-*cfmP*). **(e)** *G. mellonella* infections of wildtype, *crrB*\*L94M, *crrB*\*L94M Δ*cfmP* and *crrB*\*L94M Δ*cfmP* harboring empty vector (ev) or complementation plasmids (pACYC184-P*_lac_*::natRBS-*cfmP*). *cfmP* overexpression, complementation, and empty vector (ev) strains were grown in the presence of 500 μM IPTG.

We found that *crrB*\*L94M and CfmP induce high virulence in *G. mellonella*, causing rapid melanization and death. This severe phenotype made us wonder whether CfmP may increase host immune signaling and induce a detrimental cytokine storm. To test this, we measured how cytokine secretion and human Toll-like receptor 4 and 2 (TLR4/TLR2) signaling is impacted CrrB*L94M and CfmP. We found that wildtype *Kp* increases cytokine signaling in human macrophages (Extended Data Fig. 3a) and increases TLR4 and TLR2 activity (Extended Data Fig. 3b). However, we saw no altered immune response between wildtype and *crrB*\*L94M or *crrB*\*L94M Δ*cfmP*. Thus, while *Kp* infection stimulates the innate immune response, the virulence caused by CrrAB and CfmP is not due to abnormally high cytokine production.

### CfmP is a member of the TupA-like ATP-grasp family, with an amidoligase-like fold

*cfmP* is a gene of unknown function. Therefore, to investigate the function of CfmP, we performed protein sequence homology searches using the HHpred server^29,30^. We found that CfmP has a predicted ATP-grasp domain at its C-terminus (Fig. 3a, Extended Data Fig. 5a). ATP-grasp domains are a unique ATP-binding fold found in a large family of enzymes involved in diverse metabolic and cellular functions^31^. We performed a multi-sequence alignment of the C-terminal ATP-grasp domain of CfmP with other confirmed ATP-grasp domains and identified several conserved residues necessary for ATP binding and enzyme function (Extended Data Fig. 5a)^32,33^. To confirm that the ATP-grasp domain is necessary for CfmP’s function, we mutated the conserved E269 residue of CfmP (E269Q) to disrupt ATP-binding and found that this mutant completely reduced bacterial loads in *G. mellonella* to wildtype levels (Fig. 3b,c). 6x-Histidine tags were introduced at the C-terminus of CfmP*E269Q and wildtype CfmP, to confirm that CfmP*E269Q is stably produced (Extended Data Fig. 5b,c).

**Fig. 3:**
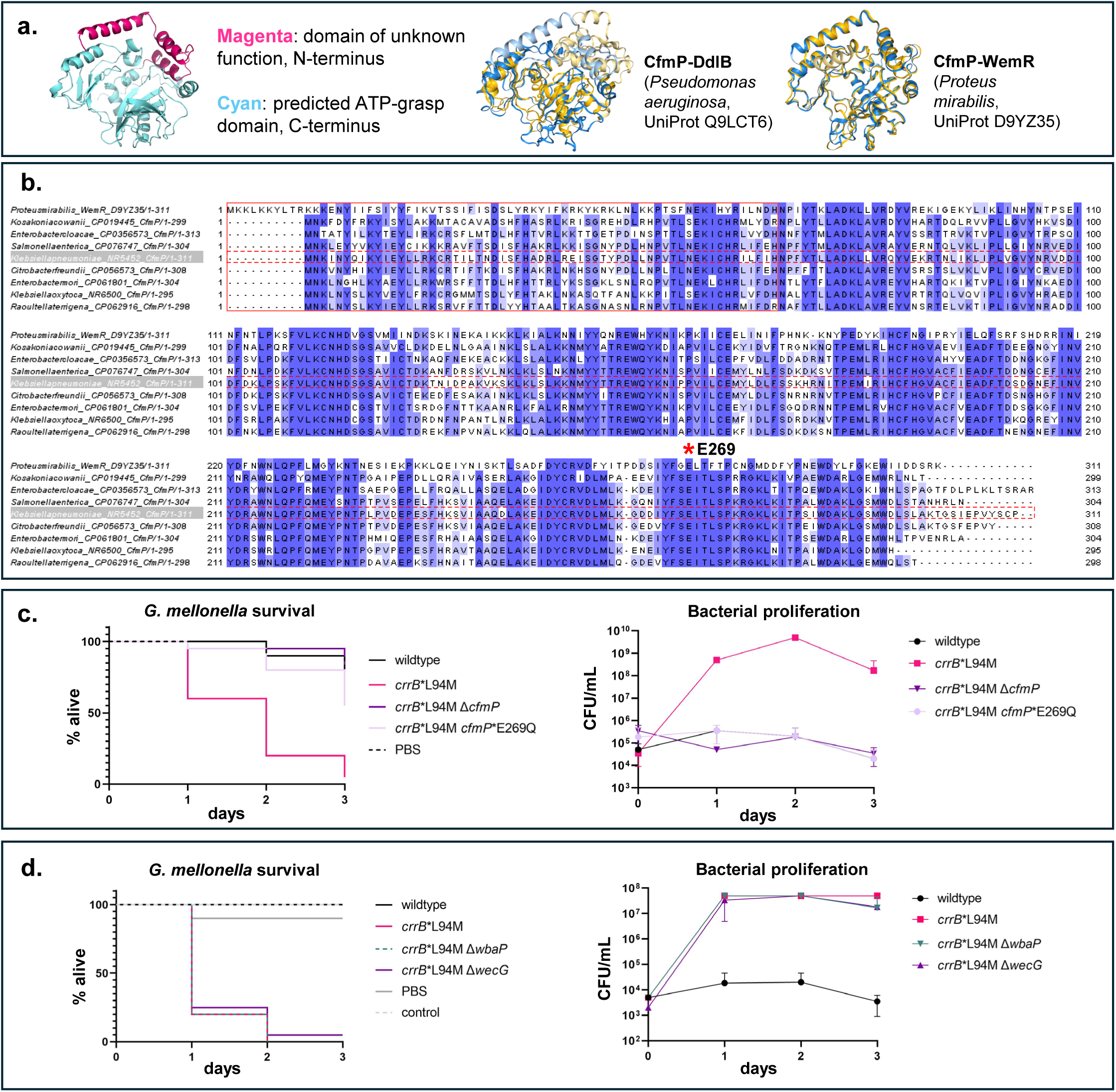
CfmP has predicted structural homology to amidoligases involved in modifying cell envelope polysaccharides. **(a)** Alphafold2-generated model of CfmP, containing a predicted ATP-grasp domain (cyan) and domain of unknown function (magenta) (left). Foldseek-generated superimpositions of the predicted structure of CfmP with DdlB from *P. aeruginosa* (UniProt Q9LCT6) (center) and with WemR from *P. mirabilis* (UniProt D9YZ35); in yellow are the structures of DdlB/WemR, in blue is the structure of CfmP. **(b)** Multiple sequence alignments of CfmP with WemR and CfmP homologs found in other *Enterobacteriaceae* species. The alignment is colored by sequence similarity, with the conserved glutamic acid (E269) starred. The N-terminal domain of unknown function of CfmP is boxed in a solid red line and the sequence of *Kp* CfmP is boxed in a dashed red line. **(c and d)** Survival curves (left) and bacterial proliferation plots (right) of *G. mellonella* infections with the indicated strains. The *crrB\**L94M Δ*wbaP* strain is deleted for the first step in capsule synthesis, and *crrB\**L94M Δ*wecG* strain is deleted for Enterobacterial common antigen (ECA).

Notably, our HHpred search did not identify any structural homology to the N-terminus of CfmP. To gain insight into the enzymatic activity of CfmP, we used AlphaFold2^34,35^ to generate a predicted structure of CfmP (Fig. 3a) and used this as an input for the Foldseek server^36^. Searches using the full sequence of CfmP clearly identified homology to ATP-grasp domains, such as that found in the D-Alanine-D-Alanine ligase DdlB (Fig. 3a), but failed to find matches to the N-terminal domain of CfmP. However, when we input only the first 61 amino acids of CfmP, we identified homology to members of the TupA-like ATP-grasp protein family (Pfam: PF14305), such as WemR from *Proteus mirabilis* (Fig. 3a)^37^. TupA-like ATP-grasp enzymes are often found within genomic loci encoding genes involved in the biosynthesis of cell surface polysaccharides and are predicted to catalyze the addition of amino acid decorations to these cell surface polymers^37,38^.

### CfmP-induced virulence does not depend on cell surface polysaccharides

Considering the predicted structural homology of CfmP to the TupA family, we tested whether the virulence induced by CfmP is dependent on the presence of cell surface polysaccharides. The cell envelope of *Kp* is composed of an inner membrane phospholipid bilayer, a structural layer of peptidoglycan, and an asymmetric outer membrane (OM) composed of an inner leaflet of phospholipids and an outer leaflet of LPS (Extended Data Fig. 6a). The OM is further decorated by the capsule and the Enterobacterial common antigen (ECA). The LPS, capsule, and ECA are all known to be immunogenic and mediate host-pathogen interactions^39,40^. Thus, we wondered whether CfmP adds a modification to one of these structures to increase virulence within the host. To test this, we individually deleted the first genes responsible for the biosynthesis of the capsule (Δ*wbaP*) and the ECA (Δ*wecG*) (Fig. 3d, Extended Data Fig. 6b) in *crrB*\*L94M. We observed no decrease in virulence (Fig. 3d), showing that CfmP must not modify these polysaccharides to induce virulence in *Kp*. Additionally, our wildtype strain of *Kp* contains an IS disrupting the O-antigen flippase and therefore produces no O-antigen (Extended Data Fig. 6b). Thus, CfmP must not affect virulence through O-antigen modifications. Finally, we performed thin layer chromatography analysis of purified lipid A from wildtype, *crrB*\*L94M, and *crrB*\*L94M Δ*cfmP* and observed no changes dependent on the presence of *cfmP* (Extended Data Fig. 6c). We performed a protein sequence alignment of WemR with *Kp* CfmP and homologs found in other Enterobacteriaceae and notably found that the N-termini of CfmP differed markedly from WemR, exhibiting only 38% percent identity (Fig. 3b). This supports that, although WemR and CfmP homologs contain similar predicted structure, their precise enzymatic activities may differ.

### Fimbriae are necessary for the virulence induced by CfmP

After ruling out the role of cell envelope polysaccharides in the virulence phenotype caused by CfmP, we returned to our RNAseq data of the highly virulent *crrB*\*L94M mutant. In addition to *cfmP*, we also found that the *fim* locus encoding type I pili, or fimbriae, is upregulated in *crrB*\*L94M (Fig. 4a,c). Fimbriae are surface exposed proteinaceous filaments important for host cell adhesion, colonization, and virulence in Gram-negative bacteria^41–43^. Fimbriae are composed of a long, helical filament of the major pilin FimA, followed by the minor pilins FimF and FimG, and the adhesin FimH at the tip of the pilus (Fig. 4a). The pilus is anchored in the OM by the usher FimD, which also aids in the export and polymerization of the FimAFGH structure^41^. Expression of the *fim* operon is known to be driven by an invertible *fimS* promoter, which under normal laboratory growth conditions is normally in the “off” orientation, keeping fimbriae production minimal^44^. However, we found that an insertion sequence (IS) is disrupting the coding region of FimE, a negative regulator of *fim* expression, in our strain background and in the original clinical isolates harboring *crrB*\*L94M (Extended Data Fig. 7a). This results in consistent production of fimbriae, even during logarithmic growth (Extended Data Fig. 7b,c).

**Fig. 4:**
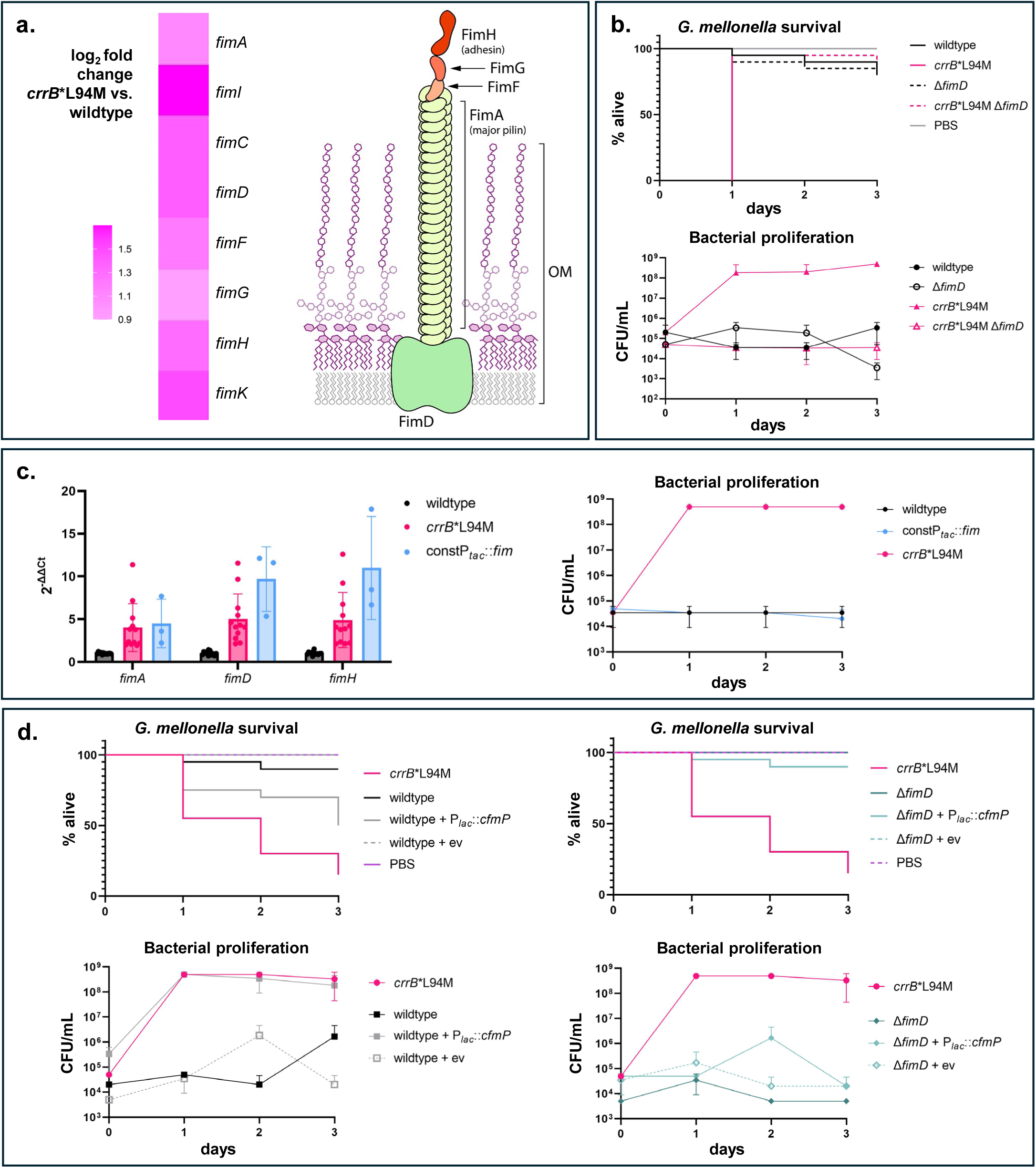
Fimbriae are essential for CfmP-driven virulence in *crrB*\*L94M. **(a)** Heat map of the log_2_ fold change of the *fim* operon in *crrB*\*L94M compared to wildtype, as identified by RNA-seq (left). Schematic of a type I pilus, or fimbriae (right). **(b)** *G. mellonella* survival curve (top) and bacterial proliferation plot (bottom) of the indicated strains. **(c)** RT-qPCR was used to confirm the increase in transcription of the *fim* operon in *crrB*\*L94M and to confirm upregulation of *fim* expression in the strain harboring an introduced optimized ribosome binding site and constitutive −10/-35 sequence upstream of *fimA* (left). *G. mellonella* bacterial proliferation curve of indicated strains (right). **(d)** *G. mellonella* survival (top) and bacterial proliferation (bottom) curves of *cfmP* overexpression strains in wildtype (left) versus Δ*fimD* (right) strains. One representative experiment is shown, split into two sets of plots for clarity. All data points are derived from the same experiment, with *crrB*\*L94M and PBS samples shown twice, on both plots, for comparison.

To investigate the role of fimbriae in *crrB*\*L94M pathogenicity, we deleted *fimD*, to completely prevent the production of pili on the surface of the cell. *G. mellonella* infections using *crrB*\*L94M Δ*fimD* found that this mutant had a virulence defect compared to *crrB*\*L94M (Fig. 4b), indicating that fimbriae are necessary for virulence in *crrB*\*L94M. Notably, deleting the usher FimD may cause a build-up of un-polymerized FimA within the cell, which could lead to fitness defects and decrease pathogenicity. To rule this out, we tested bacterial growth kinetics and found no difference in growth between *crrB*\*L94M, *crrB*\*L94M Δ*fimD*, and *crrB*\*L94M Δ*fimA* (Extended Data Fig. 7d). We wondered whether the increase in *fim* expression in *crrB*\*L94M is responsible for the higher virulence of this mutant, or whether a post-transcriptional mechanism is responsible. Thus, we introduced an optimized ribosome binding site and a constitutive −10/-35 RNA polymerase binding site from P*_tac_* upstream of the *fim* operon in the chromosome of wildtype *Kp* to artificially increase *fim* expression (Extended Data Fig. 7a). We found that this significantly increased *fim* expression but had no effects on virulence (Fig. 4c). This suggested that a post-transcriptional aspect of fimbriae production may be responsible for the pathogenicity of *crrB*\*L94M. Given the necessity of both fimbriae and CfmP for the pathogenicity of *crrB*\*L94M, we hypothesized that CfmP’s function may be to post-translationally modify fimbriae to induce virulence. To test this, we overexpressed *cfmP* in wildtype and in the absence of fimbriae and monitored bacterial loads and killing in the *G. mellonella* model. We found that the increase in virulence we saw in response to *cfmP* overexpression in wildtype did not occur in cells lacking fimbriae (Fig. 4d), indicating that fimbriae are necessary for CfmP-driven virulence.

### CfmP induces a post-translational modification to FimA to drive virulence

Together, our data indicates that CfmP is necessary for type I pili-mediated pathogenesis in *Kp*. To test whether fimbriae are post-translationally modified in a CfmP-dependent manner, we purified fimbriae from the surface of wildtype, *crrB*\*L94M, *crrB*\*L94M Δ*cfmP*, and the *cfmP*-complemented and empty vector strains and performed untargeted liquid chromatography tandem mass spectrometry (LC-MS/MS). Analyses found several peptides in FimA, FimF, and FimH containing post-translational modifications (PTMs) in *crrB*\*L94M and *crrB*\*L94M Δ*cfmP* with *cfmP* complementation (Fig. 5a, Extended Data Fig. 8). However, only one, an oxidized histidine in FimA (FimA-H11) was replicated and identified in a second biological replicate. Regardless, to test all modifications, we mutated the affected residues to alanine to prevent their modification and assessed their effects on *crrB*\*L94M virulence in *G. mellonella*. Consistent with our MS results, only mutating the oxidized histidine in FimA (FimA-H11) completely abrogated the virulence of *crrB*\*L94M (Fig. 5a,b). Interestingly, in an Alphafold^45^-generated polymer of *Kp* FimA, this histidine occurs in FimA’s N-terminal extension and points inwards towards the lumen of the polymer (Fig. 5c). Notably, the modified histidine in FimA is conserved in other *Enterobacteriaceae* species that also contain the *crr* locus (Extended Data Fig. 9b). Importantly, we confirmed that all *crrB*\*L94M *fim*\* mutants still produce fimbriae to the same levels as the parental *crrB*\*L94M strain (Extended Data Fig. 9a).

**Fig. 5:**
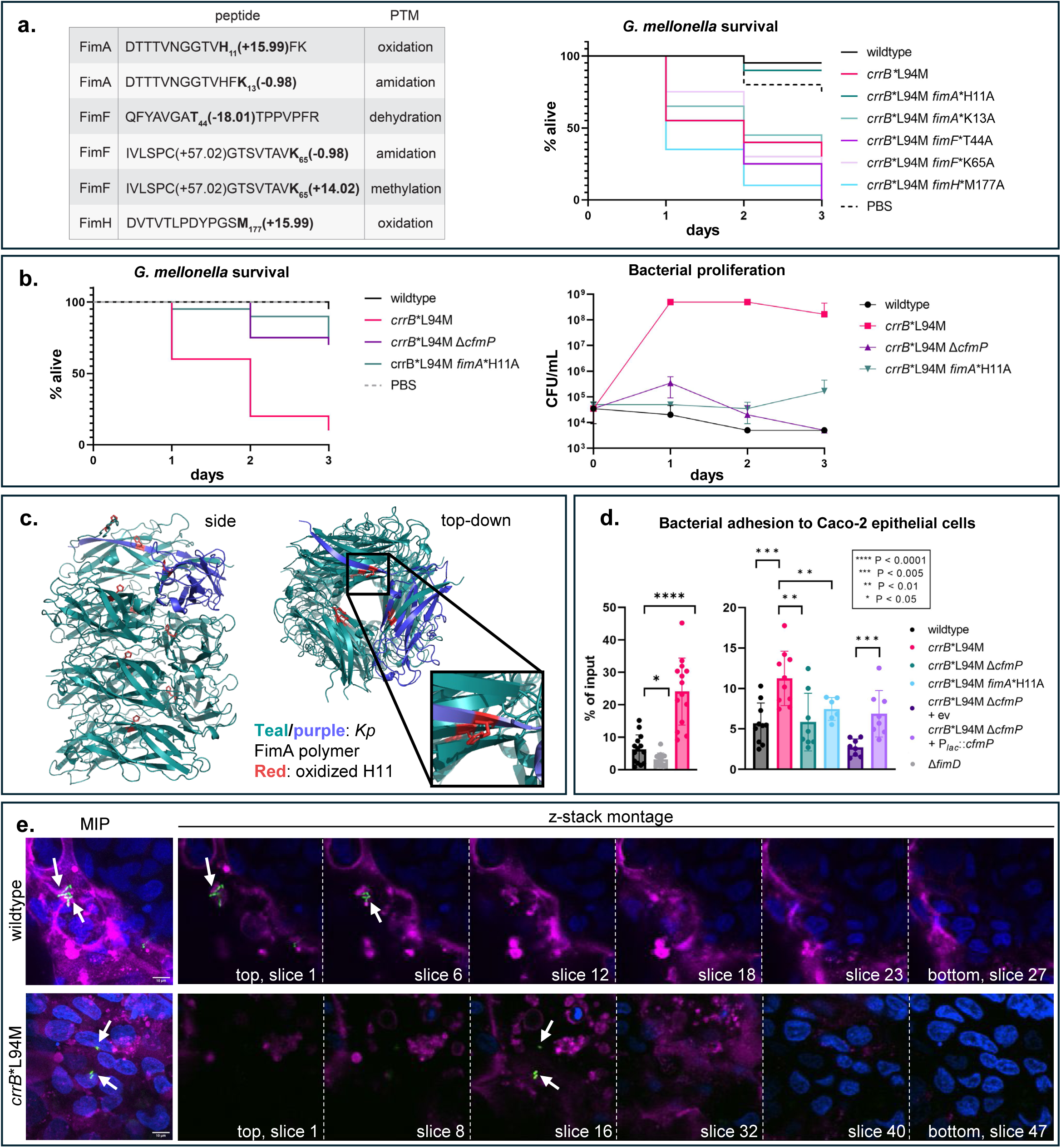
CfmP induces a post-translational modification to FimA to increase host cell adhesion and virulence. **(a)** Table of fimbriae peptides identified by untargeted LC-MS/MS to contain post-translational modifications only in *crrB*\*L94M (top). *G. mellonella* survival curve of wildtype, *crrB*\*L94M, and *crrB*\*L94M containing *fimA* and *fimF* mutants. **(b)** *G. mellonella* survival curve (left) and bacterial proliferation plot (right) of the indicated strains. **(c)** *Kp*FimA 10 subunit polymer, generated by the AlphaFold server. *Kp*FimA polymer is colored in teal, except for one subunit in purple that is highlighted. All FimA-H11 residues are shown as red sticks. **(d)** Bar charts of the indicated bacterial strains’ percent adhesion to Caco-2 monolayers after 30 minutes of incubation. Percent adhesion was calculated as the number of CFUs/mL of post-incubation bacterial suspension divided by the input CFUs/mL for each strain. **(e)** Representative confocal microscopy z-stacks of wildtype and *crrB*\*L94M adhered to Caco-2 monolayers. Bacterial strains were incubated with Caco-2 monolayers as described in the Methods. Caco-2 cells were stained for 1 h with 10 µg/mL Hoechst 33342 and for 10 m with 1X CellMask^TM^ Deep Red Plasma Membrane stain. Bacterial cells harbor a plasmid containing a Venus cytoplasmic marker. Scale bar, 10 µm. Shown are the maximum intensity projections (MIP) of all channels of each stack, and a montage of slices from each stack to show relative positions of bacteria (indicated by white arrows), Caco-2 plasma membrane (surface), and Caco-2 nuclei (interior). Each slice corresponds to 0.5 µm.

Fimbriae are important for mediating contact between pathogenic bacteria and host cells^41,42^. To test whether the CfmP-dependent oxidation on FimA-H11 impacts host-pathogen interactions, we performed adhesion assays using Caco-2 human intestinal epithelial cells^46^. We incubated monolayers of Caco-2 cells with wildtype, *crrB*\*L94M, *crrB*\*L94M Δ*cfmP*, *crrB*\*L94M *fimA*\*H11A, and Δ*fimD Kp*, washed away unadhered bacteria, plated the remaining mix of Caco-2/bacteria, and compared the number of colony forming units (CFUs) from the input bacterial cultures with the post-Caco-2 incubation output CFUs. We found that in the absence of fimbriae (Δ*fimD*) adhesion is significantly decreased compared to wildtype (Fig. 5d). Further, *crrB*\*L94M exhibits much higher adhesion than wildtype *Kp*, which is restored to wildtype levels in *crrB*\*L94M Δ*cfmP* and *crrB*\*L94M *fimA*\*H11A (Fig. 5d). Significantly, the assay we performed to test adhesion could not rule out whether output counts of *crrB*\*L94M were higher due to increased internalization of the *Kp* cells. To test this, we performed confocal microscopy of the Caco-2 cells post-incubation with *Kp*. We found that *Kp* cells were located on the surface of the Caco-2 monolayer (Fig. 5e), supporting that the CfmP-driven modification to fimbriae increases cell adhesion, and does not impact the ability of *Kp* to invade host cells.

## Discussion

Together, our data supports a model in which CrrAB induces high virulence in *Kp* through its post-transcriptional regulation of *cfmP*. We found that CfmP induces the oxidation of a highly-conserved histidine residue in the major pilin of fimbriae, FimA. CfmP-dependent modification of FimA causes increased adhesion to host cells and bacterial proliferation within the host, leading to high virulence (Fig. 6).

**Fig. 6:**
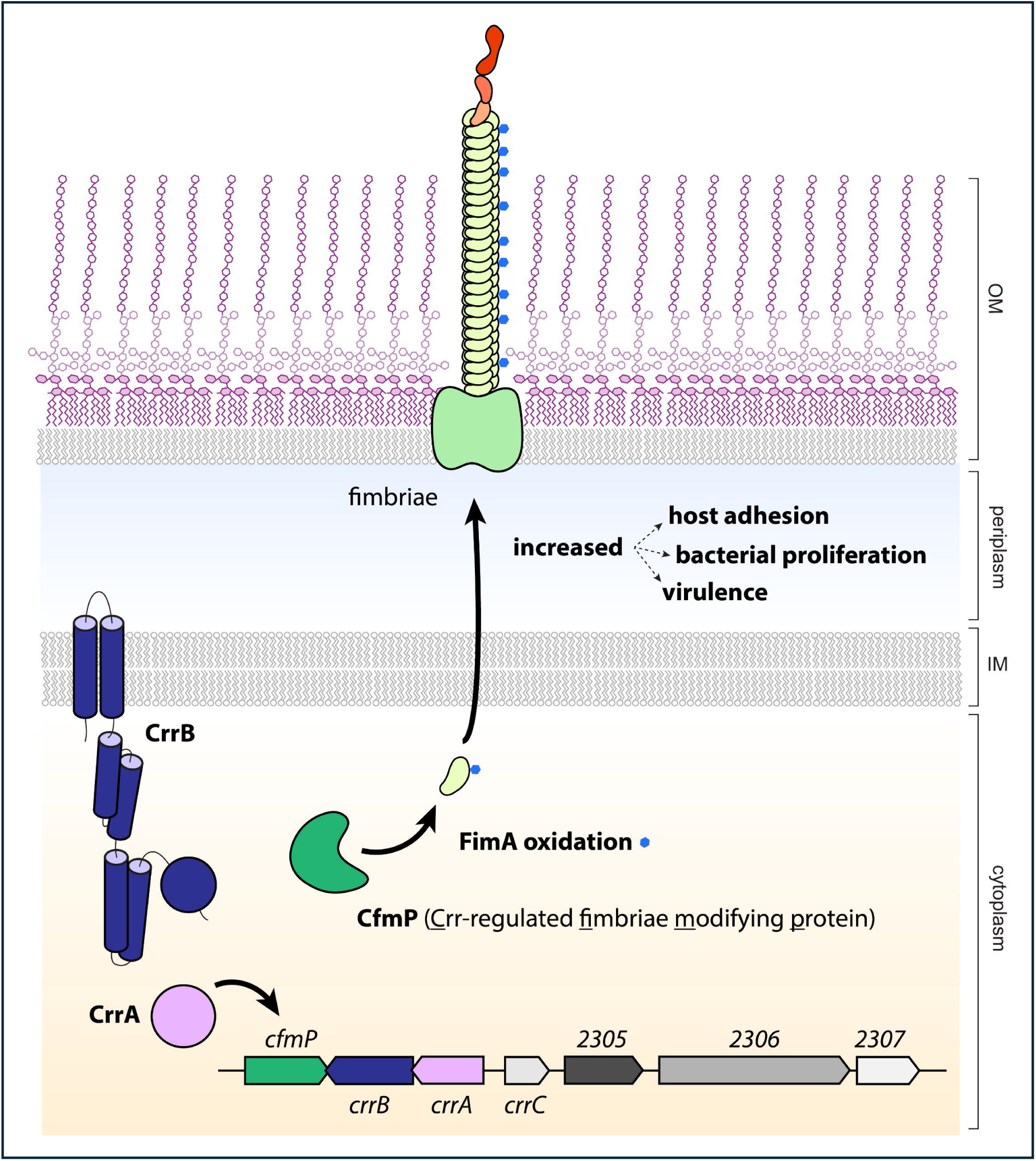
CrrAB regulation of *cfmP* induces the post-translational modification of fimbriae to increase virulence in *Kp*. The CrrAB TCS induces the expression of *cfmP* in the *crr* locus. CfmP (dark green) induces the oxidation (blue dot) of a conserved histidine in FimA (light green), the major pilin of type I pili, also known as fimbriae. The oxidized, surface-exposed fimbriae in the outer membrane cause increased adhesion to host cells. This increased adhesion then results in high bacterial proliferation in the host and a subsequent increase in virulence.

Altogether, our data is consistent with the oxidation of FimA-H11 being CrrAB- and CfmP-dependent (Fig. 5, Extended Data Fig. 8). Predicted structural homology indicates that CfmP is a novel member of the TupA-like ATP-grasp enzyme family (Fig. 3a). TupA-like ATP-grasp proteins have diverse enzymatic activities, with a subset being predicted amidoligases^31,37^. For several members of this family there is biochemical evidence supporting their amidoligase activity, such as DdlA/B and RimK^47,48^. However, the N-terminal (non-ATP-grasp) domain of CfmP differs significantly from these enzymes. Instead, CfmP’s predicted structure is more similar to WemR and other enzymes hypothesized to add amino acid modifications to OM polysaccharides (Fig. 3a)^37,38^. Despite this, our data shows that CfmP activity does not require the presence of OM polysaccharides, indicating that CfmP does not post-translationally modify any surface polysaccharide in *Kp* to induce high virulence. Consistent with this, the N-terminal domains of WemR and CfmP are not identical; many of the most highly conserved residues among CfmP homologs are not conserved in WemR (Fig. 3a,b), supporting that CfmP may have a unique function. Additionally, WemR and its homologs are often found in genomic neighborhoods with their predicted substrate, such as K- and O-antigen clusters^37,38^. *cfmP*, however, is conserved within the *crr* locus, and not found within polysaccharide biogenesis loci, further indicating that it may have a novel function. In contrast, we identified a CfmP-dependent oxidation on fimbriae, a proteinaceous extracellular polymer (Fig. 4). Our data does not rule out the possibility that the CfmP-dependent modification of fimbriae requires an additional factor to fulfill the enzymatic requirements for oxidation of FimA-H11. Biochemical analyses will be required to determine whether CfmP directly oxidizes FimA, or whether there are intermediate or interacting partners necessary for this post-translational modification.

Our data shows that CfmP increases adhesion to host cells in an intestinal epithelium model, thus leading to high bacterial loads and virulence (Fig. 5,6). Notably, we found no post-translational modifications on the adhesin FimH that impact *Kp* virulence. This suggests that CfmP does not directly affect FimH’s binding to host cells. Rather, it is a CfmP-dependent oxidation of FimA-H11 in the pilus rod that is necessary for virulence. The oxidized histidine is located within FimA’s N-terminal extension, which is critical for FimA polymerization^49^. Interestingly, the histidine is predicted to face inward towards the polymer lumen and does not obviously participate in subunit-subunit contacts^49^ (Fig. 5c). The FimA pilus is a helical polymer; the flexibility and force required to extend the helix has important implications for adhesion under shear and significantly impacts gut colonization and urinary tract infections in *E. coli*^49–53^. Thus, we hypothesize that oxidation of FimA-H11 alters the conformational dynamics of the FimA rod, allowing for better adhesion to host surfaces and subsequent colonization. Notably, *Kp* intestinal colonization is often a reservoir for systemic infections^54,55^. This suggests that fimbriae also play a role in establishing or maintaining the intestinal reservoir of *Kp* thereby increasing the risk of infection and pathogenic potential.

Genomic analyses we and others^16,17^ performed show that *crrAB* is a key regulatory system of ST258 CR*Kp*, and we hypothesized that it may be a major factor in explaining the global dominance of this important ST. Here, we confirmed that CrrAB drives the pathogenicity of ST258 through its regulation of the novel protein CfmP. In addition, we and others have shown that CrrAB increases resistance to polymyxins^16,21–23^, antibiotics with structural and functional similarities to host-produced AMPs. Significantly, we found that *crrB*\*L94M also increases resistance to the human AMP LL-37 (Extended Data Fig. 10). Thus, together our data indicates that CrrAB activity both increases resistance to host innate immunity effectors and increases the capacity of *Kp* to colonize the host environment. Together, this study supports that this TCS plays a pivotal role in the pathogenicity of *Kp* ST258.

## Online Methods

### Strains, plasmids, and routine growth conditions

Unless indicated, all *Klebsiella pneumoniae* (*Kp*) strains were derived from the parental NR5452 ST258 isolate, from the Columbia University Medical Center Clinical Microbiology Laboratory^22^. Cells were grown in Cation Adjusted Mueller Hinton broth (CAMHB) at 37°C, unless otherwise specified. Antibiotic concentrations were used at: 1 mg/mL zeocin and 50 µg/mL nourseothricin for *Kp*. Antibiotic concentrations used for cloning in DH10β were 40 µg/mL zeocin, 50 µg/mL nourseothricin, and 100 µg/mL ampicillin.

Strains used in this study can be found in Supplementary Table 1, plasmids can be found in Supplementary Table 2, and oligonucleotide primers in Supplementary Table 3.

### Strain construction

Chromosomal mutations were constructed using a CRISPR editing system^56^. Strains were electroporated with editing plasmid and plated on selective plates containing 0.2% glucose to repress Cas9/gRNA expression. Transformed colonies were streaked onto selective plates containing 0.2% arabinose to induce the CRISPR system and genome editing. Induced colonies were streaked onto non-selective media and mutations were confirmed using colony PCR. Mutants were cured of the CRISPR plasmid with serial-passaging in non-selective media.

Strains were confirmed with Sanger sequencing (Azenta). Strains used for RNAseq analysis were further confirmed by whole genome sequencing (WGS) to ensure the absence of off-target mutations. Briefly, genomic DNA was prepared using the DNeasy PowerSoil Pro Kit (Qiagen) and libraries were prepared using the Oxford Nanopore Technologies Rapid Barcoding Kit 96 V14 (SQK-RBK114.96) and sequenced using an R9 flow cell on a GridION sequencer.

### Plasmid construction

All plasmids were sequence confirmed by either Sanger sequencing (Azenta) or Oxford Nanopore sequencing (Plasmidsaurus).

**pGD13 (pUC19-CRISPR (zeoR)):** pGD13 was constructed via a 2-piece ligation. Vector backbone^56^ (Addgene #160903) was digested PvuI/AhdI and the portion containing the lambda red and *cas9* genes was retained. A PvuI/NotI-digested PCR product containing the gRNA scaffold and zeocin resistance cassette was amplified from the same plasmid with oligonucleotides oGD29/90, using splicing by overlap extension (SOE) of oGD29/88 and oGD89/90.

**pGD53 (pUC19-CRISPR (nourseR)):** pGD13 was constructed using a 2-piece isothermal assembly. pGD13 (pUC19-CRISPR (zeoR)) was digested PvuI/NotI, and the portion containing the lambda red and *cas9* genes was retained. The insert containing the gRNA scaffold and nourseothricin resistance cassette was amplified using oligonucleotides oGD29/176, from pGD13 and pGMCtN-P38-sspBmyc (Addgene #27375), using splicing by overlap extension (SOE) of oGD29/170, oKL05/06, and oGD173/176.

**pGD07 (pUC19-CRISPR-Δ*crrA*(1-150bp) (zeoR)):** pGD07 was constructed via a 2-piece ligation. Vector backbone^56^ (Addgene #160903) was digested PvuI/AhdI and the portion containing the lambda red and *cas9* genes was retained. A PCR product containing the gRNA and zeocin resistance cassette was amplified from the same plasmid with oligonucleotides oGD29/02, using splicing by overlap extension (SOE) of oGD29/12 and oGD11/02. A second PCR product containing the *crrA* deletion homology arms was amplified from NR5452 genomic DNA with oligonucleotides oGD07/08, via SOE of oGD07/04 and oGD05/08. Amplification products of oGD29/02 and oGD07/08 were further combined by SOE using oGD29/08, and digested PvuI/AhdI.

**pGD12 (pUC19-CRISPR-Δ*phoP*(1-150bp) (zeoR)):** pGD12 was constructed via a 2-piece ligation. Vector backbone^56^ (Addgene #160903) was digested PvuI/AhdI and the portion containing the lambda red and *cas9* genes was retained. A PCR product containing the gRNA and zeocin resistance cassette was amplified from the same plasmid with oligonucleotides oGD29/18, using splicing by overlap extension (SOE) of oGD29/22 and oGD21/02. A second PCR product containing the *phoP* deletion homology arms was amplified from NR5452 genomic DNA with oligonucleotides oGD17/18, via SOE of oGD17/14 and oGD15/18. Amplification products of oGD29/12 and oGD17/18 were further combined by SOE using oGD29/08, and digested PvuI/AhdI.

**pGD42 (pUC19-CRISPR-nogRNA-Δ*pmrA*(1-129bp) (zeoR)):** pGD42 was constructed using a 2-piece isothermal assembly of pGD13, digested SpeI/NotI, and a PCR product containing the *pmrA* deletion homology arms, amplified using oligonucleotides oGD189/192 via splicing by overlap extension (SOE) of PCR products oGD189/190 and oGD191/192 amplified from NR5452 genomic DNA.

**pGD47 (pUC19-CRISPR-Δ*pmrA*(1-129bp) (zeoR)):** pGD47 was constructed using a 2-piece isothermal assembly of pGD42, digested PvuI/XhoI, and a PCR product containing the gRNA amplified using oligonucleotides oGD29/63 from Addgene #160903 vector.

**pGD39 (pUC19-CRISPR-*crrB*\*L94M (zeoR)):** pGD39 was constructed using a 2-piece isothermal assembly. pGD13 was digested PvuI/AhdI, and the portion containing the lambda red and *cas9* genes was retained. A PCR product containing the gRNA, zeocin cassette, and *crrB*\*L94M homology arms was amplified from pBBR1-CRISPR-*crrB*\*L94M (zeoR)^22^ using oGD29/169.

**pGD83 (pUC19-CRISPR-nogRNA-Δ*cfmP*(1-150bp) (nourseR)):** pGD42 was constructed using a 2-piece isothermal assembly of pGD53, digested SpeI/NotI, and a PCR product containing the *cfmP* deletion homology arms, amplified using oligonucleotides oGD303/306 via splicing by overlap extension (SOE) of PCR products oGD303/304 and oGD305/306 amplified from NR5452 genomic DNA.

**pGD86 (pUC19-CRISPR-Δ*cfmP*(1-150bp) (nourseR)):** pGD86 was constructed using a 2-piece isothermal assembly of pGD83, digested PvuI/XhoI, and a PCR product containing the gRNA amplified using oligonucleotides oGD29/63 via splicing by overlap extension (SOE) of PCR products oGD29/307 and oGD308/63 amplified from pGD53.

**pGD155 (pUC19-CRISPR-nogRNA-Δ*crrC*(1-180bp) (nourseR)):** pGD155 was constructed using a 2-piece isothermal assembly of pGD53, digested SpeI/NotI, and a PCR product containing the *crrC* deletion homology arms, amplified using oligonucleotides oGD485/488 via splicing by overlap extension (SOE) of PCR products oGD485/486 and oGD487/488 amplified from NR5452 genomic DNA.

**pGD175 (pUC19-CRISPR-Δ*crrC*(1-180bp) (nourseR)):** pGD175 was constructed using a 2-piece isothermal assembly of pGD155, digested PvuI/XhoI, and a PCR product containing the gRNA amplified using oligonucleotides oGD29/63 via splicing by overlap extension (SOE) of PCR products oGD29/558 and oGD559/63 amplified from pGD53.

**pGD133 (pUC19-CRISPR-nogRNA-Δ*2305*(1-150bp) (nourseR)):** pGD133 was constructed using a 2-piece isothermal assembly of pGD53, digested SpeI/NotI, and a PCR product containing the *2305* deletion homology arms, amplified using oligonucleotides oGD103/106 via splicing by overlap extension (SOE) of PCR products oGD103/104 and oGD105/106 amplified from NR5452 genomic DNA.

**pGD136 (pUC19-CRISPR-Δ*2305*(1-150bp) (nourseR)):** pGD136 was constructed using a 2-piece isothermal assembly of pGD133, digested PvuI/XhoI, and a PCR product containing the gRNA amplified using oligonucleotides oGD29/63 via splicing by overlap extension (SOE) of PCR products oGD29/111 and oGD112/63 amplified from pGD53.

**pGD20 (pUC19-CRISPR-nogRNA-Δ*2306*(1-150bp) (nourseR)):** pGD20 was constructed using a 2-piece isothermal assembly of pGD13, digested SpeI/NotI, and a PCR product containing the *2306* deletion homology arms, amplified using oligonucleotides oGD107/110 via splicing by overlap extension (SOE) of PCR products oGD107/108 and oGD109/110 amplified from NR5452 genomic DNA.

**pGD25 (pUC19-CRISPR-Δ*2306*(1-315bp) (nourseR)):** pGD25 was constructed using a 2-piece isothermal assembly of pGD25, digested PvuI/XhoI, and a PCR product containing the gRNA amplified using oligonucleotides oGD29/63 via splicing by overlap extension (SOE) of PCR products oGD29/113 and oGD114/63 amplified from pGD53.

**pGD131 (pUC19-CRISPR-nogRNA-Δ*2306*(1-315bp) (nourseR)):** pGD131 was constructed using a 2-piece isothermal assembly of pGD53, digested SpeI/NotI, and a PCR product containing the *2306* deletion homology arms, amplified using oligonucleotides oGD107/417 via splicing by overlap extension (SOE) of PCR products oGD107/108 and oGD416/417 amplified from NR5452 genomic DNA.

**pGD135 (pUC19-CRISPR-Δ*2306*(1-315bp) (nourseR)):** pGD135 was constructed using a 2-piece isothermal assembly of pGD131, digested PvuI/XhoI, and a PCR product containing the gRNA amplified using oligonucleotides oGD29/63 via splicing by overlap extension (SOE) of PCR products oGD29/418 and oGD419/63 amplified from pGD53.

**pKL11 (pUC19-CRISPR-nogRNA-Δ*2307*(1-150bp) (nourseR)):** pKL11 was constructed using a 2-piece isothermal assembly of pGD53, digested SpeI/NotI, and a PCR product containing the *2307* deletion homology arms, amplified using oligonucleotides oKL15/18 via splicing by overlap extension (SOE) of PCR products oKL15/16 and oKL17/18 amplified from NR5452 genomic DNA.

**pKL12 (pUC19-CRISPR-Δ*2307*(1-150bp) (nourseR)):** pKL12 was constructed using a 2-piece isothermal assembly of pKL11, digested PvuI/XhoI, and a PCR product containing the gRNA amplified using oligonucleotides oGD29/63 via splicing by overlap extension (SOE) of PCR products oGD29/oKL19 and oKL20/oGD63 amplified from pGD53.

**pGD61 (pUC19 (ampR, nourseR)):** pGD61 was constructed using a 2-piece isothermal assembly of pUC19 digested AatII followed by CIP and a PCR insert containing the nourseothricin cassette amplified from pGD53 using oligonucleotides oGD226/227.

**pGD167 (pUC19-P*_lac_*::*optRBS-cfmP* (ampR, nourseR)):** pGD167 was constructed using a 2-piece isothermal assembly of pGD61 digested BamHI/PstI and a PCR insert containing *cfmP* with an optimized ribosome binding site amplified from NR5452 using oligonucleotides oGD454/547.

**pGD64 (pACYC184-P*_lac_*::MCS (tetR, catR)):** pGD64 was constructed using a 2-piece isothermal assembly of pACYC184 amplified with oGD236/oGD237 and a PCR insert containing the P*_lac_* promoter, a multi-cloning site, and a stem-loop terminator amplified from a purchased gblock (IDT) using oligonucleotides oGD238/239.

**pGD237 (pACYC184-P*_lac_*::*natRBS*-*cfmP* (tetR, catR)):** pGD237 was constructed using a 2-piece isothermal assembly of pGD64 digested SacI/PstI and a PCR insert containing *cfmP* and its native ribosome binding site amplified from NR5452 using oligonucleotides oGD797/oGD798.

**pKL01 (pUC19-CRISPR-nogRNA-crrB*M94L (nourseR)):** pKL01 was constructed using a 2-piece ligation of pGD53, digested SpeI/NotI, and a PCR product containing the *crrB*\*M94L homology arms, digested SpeI/NotI, amplified using oligonucleotides oKL01/oKL04 via splicing by overlap extension (SOE) of PCR products oKL01/oKL02 and oKL03/04, amplified from NR5452 genomic DNA.

**pKL02 (pUC19-CRISPR-crrB*M94L (nourseR)):** pKL02 was constructed using a 2-piece isothermal assembly of pKL01, digested PvuI/XhoI, and a PCR product containing the gRNA amplified using oligonucleotides oGD29/63 via splicing by overlap extension (SOE) of PCR products oGD29/thm_crrB_87_94_Up_Hom_R and thm_crrB_87_94_Down_Hom_F/oGD63, amplified from NR5452 genomic DNA.

**pGD161 (pUC19-CRISPR-nogRNA-*cfmP*\*E269Q (gRNA1) (nourseR)):** pGD161 was constructed using a 2-piece isothermal assembly of pGD53, digested SpeI/NotI, and a PCR product containing the *cfmP*\*E269Q homology arms, amplified using oligonucleotides oGD523/527 via splicing by overlap extension (SOE) of PCR products oGD523/525 and 526/527, amplified from NR5452 genomic DNA. PCR product oGD523/525 was obtained by nested PCR amplification of oGD523/524 amplified from NR5452 genomic DNA.

**pGD165 (pUC19-CRISPR-*cfmP*\*E269Q (gRNA1) (nourseR)):** pGD165 was constructed using a 2-piece isothermal assembly of pGD161, digested PvuI/XhoI, and a PCR product containing the gRNA amplified using oligonucleotides oGD29/63 via splicing by overlap extension (SOE) of PCR products oGD29/528 and oGD529/63 amplified from pGD53.

**pGD223 (pUC19-CRISPR-nogRNA-*cfmP*\*E269Q (gRNA2) (nourseR)):** pGD223 was constructed using a 2-piece isothermal assembly of pGD53, digested SpeI/NotI, and a PCR product containing the *cfmP*\*E269Q homology arms, amplified using oligonucleotides oGD523/527 via splicing by overlap extension (SOE) of PCR products oGD523/749 and oGD750/527 amplified from NR5452 genomic DNA.

**pGD225 (pUC19-CRISPR-*cfmP*\*E269Q (gRNA2) (nourseR)):** pGD225 was constructed using a 2-piece isothermal assembly of pGD223, digested PvuI/XhoI, and a PCR product containing the gRNA amplified using oligonucleotides oGD29/63 via splicing by overlap extension (SOE) of PCR products oGD29/751 and oGD752/63 amplified from pGD53.

**pGD197 (pUC19-CRISPR-nogRNA-*cfmP*-6xHis (nourseR)):** pGD197 was constructed using a 2-piece isothermal assembly of pGD53, digested SpeI/NotI, and a PCR product containing the *cfmP*-6xHis homology arms, amplified using oligonucleotides oGD666/671 via splicing by overlap extension (SOE) of PCR products oGD666/669 and 670/671, amplified from NR5452 genomic DNA. PCR product oGD666/669 was obtained by nested PCR amplification of oGD666/668 amplified from oGD666/667, amplified from NR5452 genomic DNA.

**pGD200 (pUC19-CRISPR-*cfmP*-6xHis (nourseR)):** pGD200 was constructed using a 2-piece isothermal assembly of pGD197, digested PvuI/XhoI, and a PCR product containing the gRNA amplified using oligonucleotides oGD29/63 via splicing by overlap extension (SOE) of PCR products oGD29/672 and oGD673/63 amplified from pGD53.

**pGD156 (pUC19-CRISPR-nogRNA-Δ*wbaP*(1-150bp) (nourseR)):** pGD156 was constructed using a 2-piece isothermal assembly of pGD53, digested SpeI/NotI, and a PCR product containing the *wecG* deletion homology arms, amplified using oligonucleotides oGD491/494 via splicing by overlap extension (SOE) of PCR products oGD491/492 and oGD493/494 amplified from NR5452 genomic DNA.

**pGD159 (pUC19-CRISPR-Δ*wbaP*(1-150bp) (nourseR)):** pGD159 was constructed using a 2-piece isothermal assembly of pGD156, digested PvuI/XhoI, and a PCR product containing the gRNA amplified using oligonucleotides oGD29/63 via splicing by overlap extension (SOE) of PCR products oGD29/495 and oGD496/63 amplified from pGD53.

**pGD157 (pUC19-CRISPR-nogRNA-Δ*wecG*(1-150bp) (nourseR)):** pGD157 was constructed using a 2-piece isothermal assembly of pGD53, digested SpeI/NotI, and a PCR product containing the *wecG* deletion homology arms, amplified using oligonucleotides oGD497/500 via splicing by overlap extension (SOE) of PCR products oGD497/498 and oGD499/500 amplified from NR5452 genomic DNA.

**pGD160 (pUC19-CRISPR-Δ*wecG*(1-150bp) (nourseR)):** pGD160 was constructed using a 2-piece isothermal assembly of pGD157, digested PvuI/XhoI, and a PCR product containing the gRNA amplified using oligonucleotides oGD29/63 via splicing by overlap extension (SOE) of PCR products oGD29/501 and oGD502/63 amplified from pGD53.

**pGD181 (pUC19-CRISPR-nogRNA-Δ*fimD*(1-150bp) (nourseR)):** pGD181 was constructed using a 2-piece isothermal assembly of pGD53, digested SpeI/NotI, and a PCR product containing the *fimD* deletion homology arms, amplified using oligonucleotides oGD582/585 via splicing by overlap extension (SOE) of PCR products oGD582/583 and oGD584/585 amplified from NR5452 genomic DNA.

**pGD186 (pUC19-CRISPR-Δ*fimD*(1-150bp) (nourseR)):** pGD186 was constructed using a 2-piece isothermal assembly of pGD181, digested PvuI/XhoI, and a PCR product containing the gRNA amplified using oligonucleotides oGD29/63 via splicing by overlap extension (SOE) of PCR products oGD29/586 and oGD587/63 amplified from pGD53.

**pGD228 (pUC19-CRISPR-nogRNA-Δ*fimA*(1-150bp) (nourseR)):** pGD228 was constructed using a 2-piece isothermal assembly of pGD53, digested SpeI/NotI, and a PCR product containing the *fimD* deletion homology arms, amplified using oligonucleotides oGD769/772 via splicing by overlap extension (SOE) of PCR products oGD769/770 and oGD771/772 amplified from NR5452 genomic DNA.

**pGD229 (pUC19-CRISPR-Δ*fimA*(1-150bp) (nourseR)):** pGD229 was constructed using a 2-piece isothermal assembly of pGD228, digested PvuI/XhoI, and a PCR product containing the gRNA amplified using oligonucleotides oGD29/63 via splicing by overlap extension (SOE) of PCR products oGD29/773 and oGD774/63 amplified from pGD53.

**pGD231 (pUC19-CRISPR-nogRNA-constP*_tac_*::*fim* (nourseR)):** pGD231 was constructed using a 2-piece isothermal assembly of pGD53, digested SpeI/NotI, and a PCR product containing the *fimD* deletion homology arms, amplified using oligonucleotides oGD782/785 via splicing by overlap extension (SOE) of PCR products oGD782/783 and oGD784/785 amplified from NR5452 genomic DNA.

**pGD233 (pUC19-CRISPR-constP*_tac_*::*fim* (nourseR)):** pGD233 was constructed using a 2-piece isothermal assembly of pGD231, digested PvuI/XhoI, and a PCR product containing the gRNA amplified using oligonucleotides oGD29/63 via splicing by overlap extension (SOE) of PCR products oGD29/786 and oGD787/63 amplified from pGD53.

**pGD238 (pUC19-CRISPR-nogRNA-*fimA*\*H11A (nourseR)):** pGD238 was constructed using a 2-piece isothermal assembly of pGD53, digested SpeI/NotI, and a PCR product containing the *fimA*\*H11A mutant homology arms, amplified using oligonucleotides oGD800/801 via splicing by overlap extension (SOE) of PCR products oGD800/804 and oGD805/801 amplified from NR5452 genomic DNA.

**pGD240 (pUC19-CRISPR-*fimA*\*H11A (nourseR)):** pGD240 was constructed using a 2-piece isothermal assembly of pGD238, digested PvuI/XhoI, and a PCR product containing the gRNA amplified using oligonucleotides oGD29/63 via splicing by overlap extension (SOE) of PCR products oGD29/817 and oGD818/63 amplified from pGD53.

**pGD239 (pUC19-CRISPR-nogRNA-*fimA*\*K13A (nourseR)):** pGD239 was constructed using a 2-piece isothermal assembly of pGD53, digested SpeI/NotI, and a PCR product containing the *fimA*\*K13A mutant homology arms, amplified using oligonucleotides oGD800/801 via splicing by overlap extension (SOE) of PCR products oGD800/806 and oGD807/801 amplified from NR5452 genomic DNA.

**pGD241 (pUC19-CRISPR-*fimA*\*K13A (nourseR)):** pGD241 was constructed using a 2-piece isothermal assembly of pGD239, digested PvuI/XhoI, and a PCR product containing the gRNA amplified using oligonucleotides oGD29/63 via splicing by overlap extension (SOE) of PCR products oGD29/817 and oGD818/63 amplified from pGD53.

**pGD242 (pUC19-CRISPR-nogRNA-*fimF*\*T44A (nourseR)):** pGD242 was constructed using a 2-piece isothermal assembly of pGD53, digested SpeI/NotI, and a PCR product containing the *fimF*\*T44A mutant homology arms, amplified using oligonucleotides oGD808/809 via splicing by overlap extension (SOE) of PCR products oGD808/810 and oGD811/809 amplified from NR5452 genomic DNA.

**pGD245 (pUC19-CRISPR-*fimF*\*T44A (nourseR)):** pGD245 was constructed using a 2-piece isothermal assembly of pGD242, digested PvuI/XhoI, and a PCR product containing the gRNA amplified using oligonucleotides oGD29/63 via splicing by overlap extension (SOE) of PCR products oGD29/819 and oGD820/63 amplified from pGD53.

**pGD243 (pUC19-CRISPR-nogRNA-*fimF*\*K65A (nourseR)):** pGD243 was constructed using a 2-piece isothermal assembly of pGD53, digested SpeI/NotI, and a PCR product containing the *fimF*\*K65A mutant homology arms, amplified using oligonucleotides oGD808/809 via splicing by overlap extension (SOE) of PCR products oGD808/812 and oGD813/809 amplified from NR5452 genomic DNA.

**pGD246 (pUC19-CRISPR-*fimF*\*K65A (nourseR)):** pGD246 was constructed using a 2-piece isothermal assembly of pGD243, digested PvuI/XhoI, and a PCR product containing the gRNA amplified using oligonucleotides oGD29/63 via splicing by overlap extension (SOE) of PCR products oGD29/821 and oGD822/63 amplified from pGD53.

**pGD244 (pUC19-CRISPR-nogRNA-*fimH*\*M177A (nourseR)):** pGD243 was constructed using a 2-piece isothermal assembly of pGD53, digested SpeI/NotI, and a PCR product containing the *fimH*\*M177A mutant homology arms, amplified using oligonucleotides oGD828/829 via splicing by overlap extension (SOE) of PCR products oGD828/830 and oGD831/829 amplified from NR5452 genomic DNA.

**pGD247 (pUC19-CRISPR-*fimH*\*M177A (nourseR)):** pGD247 was constructed using a 2-piece isothermal assembly of pGD244, digested PvuI/XhoI, and a PCR product containing the gRNA amplified using oligonucleotides oGD29/63 via splicing by overlap extension (SOE) of PCR products oGD29/832 and oGD833/63 amplified from pGD53.

**pGD63 (pACYC184-empty-MCS (catR, tetR)):** pGD63 was constructed using a 2-piece isothermal assembly of pACYC184 amplified with oGD247/248, and a PCR product containing a multi-cloning site and Rho-independent terminator amplified from pGD15^57^ with a nested PCR (first oGD249/250, second oGD249/251).

**pGD70 (pACYC184-optRBS-*venus* (catR, tetR)):** pGD70 was constructed using a 2-piece ligation of pGD63, digested SacI/PstI, and a PCR product containing venus and an optimized ribosome binding site amplified from pGD15^57^ using oGD272/273, digested SacI/PstI.

**pGD91 (pACYC184-P*_tac_*::optRBS-*venus* (catR, tetR)):** pGD91 was constructed using a 2-piece isothermal assembly of pGD70 digested SpeI/SacI, and an insert containing the −10 and - 35 sequence from P*_tac_*, without the operator, created by hybridizing oligos oGD329/330.

### Sequence typing and *crrAB* genomic searches

Previously-sequenced carbapenem resistant *Kp* isolates collected from Columbia University Irving Medical Center from 2009-2023 were used for genomic analyses^23–25^. Multi-locus sequence typing (MLST) and genomic searches for *crrAB* were performed using SRST2^58^. For *crrAB* searches, the first ST258 *Kp* isolate collected per patient was included in the analysis.

### Gene ontology analysis

Genomic neighborhood analysis was performed using the Enzyme Function Initiative Genome Neighborhood Tool5 (EFI-GNT v1.0) available at: https://efi.igb.illinois.edu//efi-gnt/index.php. The sequence of CrrB was used as the query for a BLAST search of the UniProt database, using the "Retrieve neighborhood diagrams" option within EFI-GNT. Gene neighborhood diagrams (GND) were generated to visualize the 20 nearest genes surrounding *crrB*.

### RNAseq

Bacterial strains were grown to mid-log phase in Cation-adjusted Mueller Hinton Broth (CAMHB) at 37°C and cells were harvested and normalized to 500 μL of OD_600_ = 0.5. Cell pellets were resuspended in RNAprotect (Qiagen) and RNA was extracted using the RNeasy kit with an on-column DNase treatment (Qiagen). RNAseq libraries were prepared using the QIAseq UPXome RNA Library Kit for Bacteria, with FastSelect rRNA depletion (Qiagen). Sequencing was performed on an Illumina MiSeq using a v2 300 cycle reagent kit. Quality-filtered reads (*trimmomatic*^59^) were mapped to the NR5452 wildtype single contig reference genome using *bowtie2*^60^, and *samtools*^61^ and *htseq*^62^ were used to generate read count tables. *DESeq2*^63^ in R was used to identify genes with significantly altered transcription. Heatmaps were generated in R using ggplot2^64^.

### Quantitative reverse transcription PCR (RT-qPCR)

Bacterial strains were grown to mid-log phase in Cation-adjusted Mueller Hinton Broth (CAMHB) at 37°C and cells were harvested and normalized to 500 μL of OD_600_ = 0.5. Cell pellets were resuspended in RNAprotect (Qiagen) and RNA was extracted using the RNeasy kit with an on-column DNase treatment (Qiagen). cDNA was generated from 500 ng input RNA using LunaScript RT SuperMix Kit (NEB) and was used as a template for RT-qPCR reactions using SsoAdvanced Universal Inhibitor-Tolerant SYBR Green Supermix (BioRad). Log fold changes were calculated using the 2^−ΔΔC^_T_ method^65^, with *rpsL* used as the housekeeping gene.

### Drug susceptibility testing

Minimum inhibitory concentrations (MICs) were determined using broth microdilution (BMD) according to Clinical and Laboratory Standards Institute guidelines (CLSI, 2017). Suspensions of each bacterial strain were made to a MacFarland standard of 0.5, diluted 1:100 in Cation-Adjusted Mueller-Hinton broth (CAMHB), and inoculated into a 96 well plate containing increasing concentrations of antibiotic/compound in CAMHB and incubated for 24 hours at 37°C.

### *Galleria mellonella* infections

Bacterial strains were grown in Cation-adjusted Mueller Hinton Broth (CAMHB) at 37°C to mid-log phase, and cells were harvested and normalized to OD_600_ of 0.05 in phosphate buffered saline (PBS). 10 μL of each bacterial suspension (corresponding to 1 x 10^7^ colony forming units (CFUs)/mL, or 10^5^ CFUs total per larva) was injected into the last left proleg of 20 *G. mellonella* larvae per strain^66^. 20 larvae were injected with 10 μL of PBS as a negative control. For experiments in which larvae were additionally plated out for bacterial loads, 20 additional larvae were injected per strain for a total of 40 larvae per strain. Serial dilutions of each strain suspension were plated to calculate CFUs and ensure equivalent inoculums. Larvae mortality was monitored over the course of 3 to 7 days by tactile stimulus^66^. To calculate bacterial proliferation, 3 larvae per strain, per day were each homogenized in 500 μL PBS, and serial dilutions were plated on KPC Chromagar^TM^ agar plates to select for the injected carbapenem resistant *Kp*. Larvae injected with only PBS showed no growth on KPC plates. 2 μL of 10^-1^ to 10^-6^ serial dilutions of homogenized *G. mellonella* were plated and dilution at which growth stopped was used calculate total bacterial load per larva.

### Cytokine production panel

THP-1 monocytes were obtained from ATCC (TIB-202). Cells were maintained in ATCC-formulated RPMI-1640 medium containing 10% fetal bovine serum (FBS), 100 U/mL penicillin, and 100 μg/mL streptomycin at 37°C with 5% CO_2_. THP-1 cells were seeded into 12-well culture-treated plates at 5×10^5^ cells per well and were incubated with 200 nM phorbol 12-myristate 13-acetate (PMA) for 72 hours to differentiate into macrophages. 24 hours prior to infection/incubation with antigen, differentiated and adhered THP-1 cells were washed twice with phosphate buffered saline (PBS) and wells were replenished with fresh media, without PMA.

Bacterial strains were grown to mid-log phase and 1.5 mL of OD_600_ = 0.05 was harvested and resuspended to concentrate in 60 μL of PBS. 20 μL of bacteria were added to each well of differentiated THP-1 macrophages, for an approximate MOI = 10. As a positive control, purified LPS from *Escherichia coli* 0111:B4 (InvivoGen) was added to a final concentration of 100 ng/mL. Plates were then incubated at 37°C with 5% CO_2_ for 4 hours. Input bacterial suspensions were serially diluted and plated for CFUs to confirm equal bacterial loads between strains. After 4 hours, supernatant containing secreted cytokines was collected from wells, filter sterilized (0.22 μm), and stored at −80°C. Samples were analyzed using the Human Cytokine Proinflammatory Focused 15-Plex Discovery Assay Array (HDF15) from Eve Technologies.

### TLR4/TLR2 signaling assays

HEK-Blue^TM^ hTLR4 and hTLR2 cells were obtained from InvivoGen. Cells were maintained in Dulbecco’s Modified Eagle Medium (DMEM) containing 10% fetal bovine serum (FBS), 100 μg/mL Normocin^TM^, 100 U/mL penicillin, 100 μg/mL streptomycin, and HEK-Blue Selection^TM^ at 37°C with 5% CO_2_.

For signaling assay, HEK-Blue cells were suspended in HEK-Blue^TM^ Detection medium at 140,000 cells/mL and 180 μL was added per well (corresponding to 25,000 cells) into 96-well culture-treated plates. Bacterial strains were grown to mid-log phase and normalized to OD_600_ = 0.03125 in PBS (corresponding to a multiplicity of infection of 5 in 20 μL). Bacteria were heat killed at 80°C for 20 minutes and serially diluted to the appropriate MOI in equivalent volumes (ranging from 0.01 to 5). Plates were incubated at 37°C with 5% CO_2_ for 18 hours. Plates were briefly shaken, and OD_640_ measurements were taken.

### Sequence Alignment

Protein sequences of ATP-grasp domain containing proteins and FimA were obtained from the UniProt KnowledgeBase^67^ (UniProtKB; https://www.uniprot.org/) or in-house isolated and sequenced clinical strains. Multi-sequence alignments were performed using either MUSCLE or Clustal Omega^68^. Pairwise sequence alignments were performed and percent identity calculated using EMBOSS Needle^68^. Alignments were visualized in Jalview^69^.

### Immunobloting

The OD_600_ was recorded for each culture, and cells were harvested to a final, normalized OD_600_ of 10 for equivalent loading. Pellets were collected and re-suspended in 1X sodium dodecyl sulfate (SDS) sample buffer (10% glycerol, 50mM TrisHCl pH 6.8, 2% SDS, 0.005% bromophenol blue) containing 10% β-mercaptoethanol. Cells were heated at 95°C for 10 min prior to loading. Proteins were separated by SDS-PAGE on 4-20% TGX polyacrylamide gels (Biorad), transferred onto nitrocellulose membranes (Biorad), and blocked in 3% bovine serum albumin (BSA) in phosphate-buffered saline (PBS) with 0.5% Tween-20. The blocked membranes were probed with anti-His (1:5,000) (GenScript) or anti-FimA (1:5,000) (Invitrogen) diluted into 3% BSA in 1x PBS with 0.05% Tween-20. Primary antibodies were detected using horseradish peroxidase-conjugated goat anti-mouse or anti-rabbit IgG (BioRad) and the Clarity™ Western ECL Substrate (Biorad). Signal was detected using a ChemiDoc MP Imaging System (Biorad).

### Cell envelope polysaccharide detection

Bacterial strains were grown to mid-log phase and cells were harvested and normalized to 500 μL of OD_600_ = 1.0. Cell pellets were resuspended in 50 μL 1x SDS sample buffer (10% glycerol, 50mM TrisHCl pH 6.8, 2% SDS, 0.005% bromophenol blue) containing 10% β-mercaptoethanol. Samples were boiled 10 minutes, then incubated overnight at 55°C with 300 μg/mL Proteinase K (Qiagen). Samples were run at 150V for 1 hour on a 4-12% Bis-Tris polyacrylamide gel (Biorad) and capsule / LPS was labeled using the Pro-Q™ Emerald 300 Glycoprotein Gel and Blot Stain Kit (Invitrogen) following the manufacturer’s instructions.

### Lipid A radiolabeling and thin layer chromatography

To evaluate any changes in the lipid A domain of LPS, *E. coli* and *K. pneumoniae* were grown in the presence of 2.5 μCi/ml of ³²P_i_ (Perkin-Elmer). The lipid portion of LPS was released using mild acid hydrolysis (pH 4.5, 100°C, 30 min), which cleaves the bond between lipid A and the first residue of the core oligosaccharide. Lipid A species were then isolated following established protocols^70,71^ and resolved by thin-layer chromatography (TLC) using a solvent mixture of chloroform, pyridine, 88% formic acid, and water (50:50:16:5, vol/vol). Radiolabeled species were detected by phosphorimager analysis. *E. coli* K-12 strains W3110 and WD101 were included as controls. W3110 produces unmodified lipid A and is polymyxin sensitive, whereas the lipid A of WD101 is highly modified and is polymyxin resistant.

### Fimbriae purification

Bacterial strains were grown to late-log phase in CAMHB at 37°C, and 125 mL of OD_600_ of 1.0 was harvested as normalized input. Cells were pelleted at 5000 xg for 30 minutes and washed twice in 2.5 mL 10 mM Tris pH 8. Suspensions were incubated at 65°C for 1 hour to release fimbriae from the cell surface. Cells were pelleted and supernatant containing fimbriae was retained. NaCl was added to 500 mM final and MgCl_2_ was added to 250 mM final, and supernatants were incubated with rocking at 4°C overnight. Precipitated protein was spun at 14,000 xg, high salt buffer was removed, and fimbriae were resuspended in 10 mM Tris pH 8 to re-solubilize proteins. Protein concentration was measured using a BCA Protein Assay Kit (Pierce). Equal volumes of purified fimbriae from each strain were run on 4-12% Bis-Tris polyacrylamide gel (Biorad) and stained with GelCode Blue Stain Reagent (ThermoFisher).

### Transmission electron microscopy (TEM)

Bacterial strains were grown to log phase in CAMHB at 37°C. Cultures were concentrated to 100 μL of OD_600_=10 and resuspended in 500 μL of 2% glutaraldehyde in 0.1X PBS for fixation. Cells were fixed for 1 hour at 4°C and then washed 5 times in 500 μL 0.1X PBS. Cell suspensions were spotted onto 200 copper mesh grids coated in pure carbon (Ted Pella) and imaged on a FEI Talos F200X Transmission / Scanning Electron Microscope.

### Mass spectrometry and data analysis

Mass spectrometry analyses were performed at the Mass Spectrometry Facility, UMass Chan Medical School. Protein samples were digested overnight at 37°C with trypsin (Promega) using the S-Trap method. Peptides were eluted with 100 µL of 50% acetonitrile in water, dried in a SpeedVac, and reconstituted in 20 µL of 0.1% formic acid in 5% acetonitrile. The reconstituted samples were centrifuged at 16,000×g for 16 min, and 18 µL of the supernatant was transferred to low-binding HPLC vials. A 1 µL aliquot was injected into a TimsTOF Pro2 mass spectrometer (Bruker) coupled to a nanoElute LC system (Bruker).

Chromatographic separation was performed on an in-house–packed C18 column (250 mm × 75 µm i.d., 3 µm particle size, 120 Å pore size) using a 30 min linear gradient at a flow rate of 500 nL/min. The mobile phases consisted of Solvent A (0.1% formic acid in water) and Solvent B (0.1% formic acid in acetonitrile).

Data were acquired in Data-Dependent Acquisition mode using Parallel Accumulation–Serial Fragmentation (DDA-PASEF). Each acquisition cycle comprised 10 PASEF MS/MS scans, with 100 ms ramp and accumulation times. The m/z range was set to 100–1700, and the ion mobility range (1/K₀) to 0.70–1.30 V s/cm². Collision energy was linearly scaled with ion mobility, from 20 eV at 0.6 V s/cm² to 59 eV at 1.6 V s/cm². Instrument calibration was performed using the ESI-L Tuning Mix (Agilent) with reference ions at m/z 622, 922, and 1222.

Raw data were processed using two independent workflows for confirmation. First, FragPipe/MSFragger was used in a non-specific search mode against a custom protein database (listed below), combined to a contaminant database, to identify peptides in each sample, including those with post-translational modifications. Results from FragPipe/MSFragger were visualized in Scaffold (version 5.2.2, Proteome Software Inc.). Second, the +15.99 Da modification on the FimA histidine residue in the peptides identified in Scaffold was selected for targeted validation in Skyline (64-bit, version 25.1.0.142), using monoisotopic precursor and fragmented masses including precursor/ion charge states from 1 to 4.

Custom protein database:

- *FimA_FloresKim_Dobihal* MKIKTLAMIVVSALSLSSTAALADTTTVNGGTVHFKGEVVNAACAVDAGSIDQTVQLGQVRSAKLATAGSTSSAVGFNIQLDDCDTTVATKASVAFAGTAIDSSNTTVLALQNSAAGSATNVGVQILDNTGTPLALDGATFSAATTLNDGPNIIPFQARYYATGAATAGIANADATFKVQ YE
- *FimF_FloresKim_Dobihal* MRTLQYLLGALFTLGAPAALAADSTIAISGYVRDNACTVAGESKDFTVDLQDNAAKQFYAVGATTPPVPFRIVLSPCGTSVTAVKVGFTGVADSVNTSLLKLDAGTSAAAGMGVEILDQQQSRLPVNAPSSTMTWTTLTPGQTNILNFYARLMATQVPVTAGHVNATATFTLEFQ
- *FimG_FloresKim_Dobihal* MKWTHVGWLLAGLLTASASLRAADVTLTVNGKVVARPCTVSTVNATVELGDLYTFSLIGAGSASAWHSVALDLSNCPVGTSRVKATFSGTADSTGYYKNQGTAGNIQLELQNEDGTTLNNGSSQSVQVDEASQSARFPLQVRALSVNGGATQGTIQAVINVTYTYA
- *FimH_FloresKim_Dobihal* MMKKIIPLFTTLLLLGWSMNAWSFACKTATGATIPIGGGSANVYVNLTPAVNVGQNLVVDLSTQIFCHNDYPETITDYVTLQRGSAYGGVLSSFSGTVKYNGTSYPFPTTTETARVIYDSRTDKPWPAVLYLTPVSTAGGVAITAGSLIAVLILHQTNNYNSDSFQFIWNIYANNDVVVPTGGCDVSARDVTVTLPDYPGSMAVPLTVHCAQSQQLGYYLSGTTADSANAIFTNTASASPAQGIGVQLTRNGSAVPANSTVSLGTVGTSPVNLGLTATYARTTGQVTAGNVQSIIGITFV YQ

### Epithelial cell adhesion assay

Caco-2 colon epithelial cells were obtained from ATCC (HTB-37). Cells were maintained in ATCC-formulated Eagle’s Minimum Essential Medium (EMEM) containing 20% fetal bovine serum (FBS), 100 U/mL penicillin, and 100 μg/mL streptomycin at 37°C with 5% CO_2_. To prepare cells for the bacterial adhesion assay, Caco-2 cells were seeded at 2.1×10^5^ cells/mL per well in a culture-treated 12-well plate. Caco-2 layers were differentiated for 14 days, with media renewed every three days. Prior to adding bacteria, old media was removed from the Caco2 cells, and fresh media without antibiotics was added to each well.

Bacterial strains were grown to mid-log phase and normalized to OD_600_ = 0.25 in 500 μL of PBS. 20 μL of bacteria were added to each well of Caco-2 cells (for an approximate MOI = 5) and were incubated at 37°C for 30 minutes. Input bacterial suspensions were serially diluted and plated for CFUs. After incubation, wells were washed 5 times with 500 μL of PBS, and contents were resuspended in 1 mL ice-cold 0.01% Triton X-100. Suspensions were serially diluted and plated for output CFUs. Adhesion/internalization was calculated as the % remaining of total input bacterial counts, post-washes for each strain.

### Confocal microscopy

Caco-2 cells were cultured as described for the “Epithelial cell adhesion assay” but were grown in a 1.5 uncoated coverslip 10 mm diameter 12-well plate (MatTek). 1 hour prior to infections, 10 μg/mL Hoechst 33342 (BD Biosciences) was added to the cultures. Bacterial strains were grown to mid-log and normalized to OD_600_ = 0.25 in 500 μL of PBS. 20 μL of bacteria were added to each well of Caco-2 cells (for an approximate MOI = 5) and were incubated at 37°C for 30 minutes. Input bacterial suspensions were serially diluted and plated for CFUs. After incubation, wells were washed 2 times with 500 μL of PBS, then stained for 10 minutes with 5 μg/mL CellMask™ Deep Red Plasma Membrane Stains (Invitrogen) in 1X PBS. Wells were washed another 3 times with 500 μL of PBS prior to imaging.

Confocal images were acquired using an AXR confocal scanner mounted on a Nikon Ti2 inverted microscope equipped with a 60x/1.49 ApoTIRF oil-immersion objective (Nikon). Images were acquired using gallium arsenide phosphide (GaAsP) (green and red fluorophores) or conventional silicon (blue and far-red fluorophores) PMTs, sequentially, using resonant scanning at a pixel size of 0.14 μm/px and scan zoom of 1.00x. Z-stacks were collected at intervals of 0.5 µm using the internal microscope Z drive; total depth per z-stack was determined empirically for each field of view, to maximize capturing the top of the sample and the interior.

Venus was illuminated at 488 nm with power 17.4 and gain 46.7 and emission was collected from 517-545 nm; Hoechst 33342 was illuminated at 405 nm with power 20.0 and gain 36.7 and emission was collected from 429-474 nm; CellMask^TM^ Deep Red was illuminated at 640 nm with power 4.7 and gain 15.1 and emission was collected from 660-750 nm. At least five fields of view were acquired for each sample. Images were acquired using Nikon Elements 4.3 acquisition software. Image processing was performed using Fiji and the 3DViewer plugin.

### Statistics and reproducibility

All experiments were performed in at least biological triplicate and representative images/results or total data points with standard deviation and mean are shown. Attempts at replication for all experiments was successful.

## Acknowledgments

We thank all members of the Uhlemann and Dworkin labs for discussions and encouragement. We also thank Ed Lopatto and Henry Schreiber of the Hultgren lab for helpful advice and insights on working with fimbriae.

This study used the Confocal and Specialized Microscopy Shared Resource of the Herbert Irving Comprehensive Cancer Center at Columbia University, funded in part through the NIH/NCI Cancer Center Support Grant P30CA013696. This study additionally used the Mass Spectrometry Facility at UMass Chan Medical School. We acknowledge them for their technical expertise and instrumentation for proteomics analyses.

This work was funded by NIH grants T32AI100852 (G.S.D.), R01AI174416 (M.S.T.), R01AI176776 (M.S.T.), R35GM150669 (J.F.K.), Smith Family Awards Program for Excellence in Biomedical Research (J.F.K.), K24AI183182 (A.C.U.), and U54DK104309 (A.C.U.).

## Author contributions

G.S.D., J.F.K., and A.C.U. conceived the study. G.S.D., K.L., L.Y., and C.M.H. performed the experiments. G.S.D., K.L., J.F.K., C.M.H., M.S.T., and A.C.U. analyzed data. G.S.D. wrote the original draft of the paper with revisions and edits contributed by J.F.K., M.S.T., and A.C.U.

---

**Extended Data Fig. 1:**
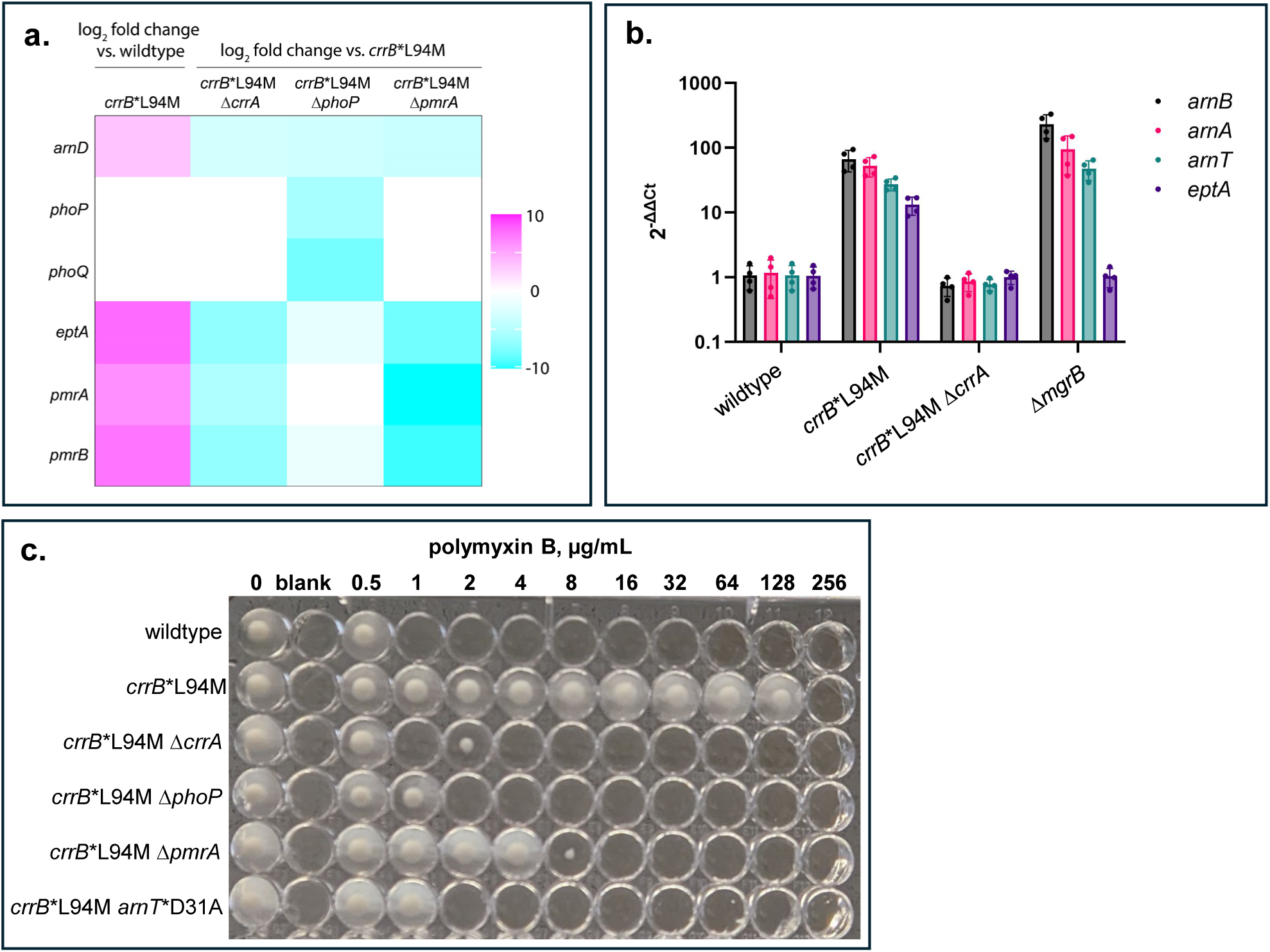
CrrAB regulates the *arn* operon and *pmrAB-eptA*. **(a)** Heat map of the log_2_ fold change of *arnD, phoP, phoQ, eptA, pmrA*, and *pmrB* transcripts in the indicated strains as identified by RNA-seq. Log_2_ fold change is shown for one gene in the *arn* operon, *arnD,* encoding the final enzyme in the amino arabinose synthesis pathway before it is flipped to the cytoplasm and ligated to lipid A. **(b)** RT-qPCR was used to confirm the change in transcription of *arnB*, *arnA*, *arnT*, and *eptA* in the indicated strains. A deletion of *mgrB* (Δ*mgrB*), encoding a negative regulator of PhoPQ, was deleted, was included to compare activation levels of the indicated genes through CrrAB vs. PhoPQ signaling. **(c)** Polymyxin B broth microdilution (BMD) plate of the indicated mutants. A catalytically inactive mutant of ArnT (*arnT*\*D31A) (Petrou *et al.* 2016) also decreased polymyxin resistance in *crrB*\*L94M, confirming that induction of the *arn* operon is necessary for resistance in this background.

**Extended Data Fig. 2:**
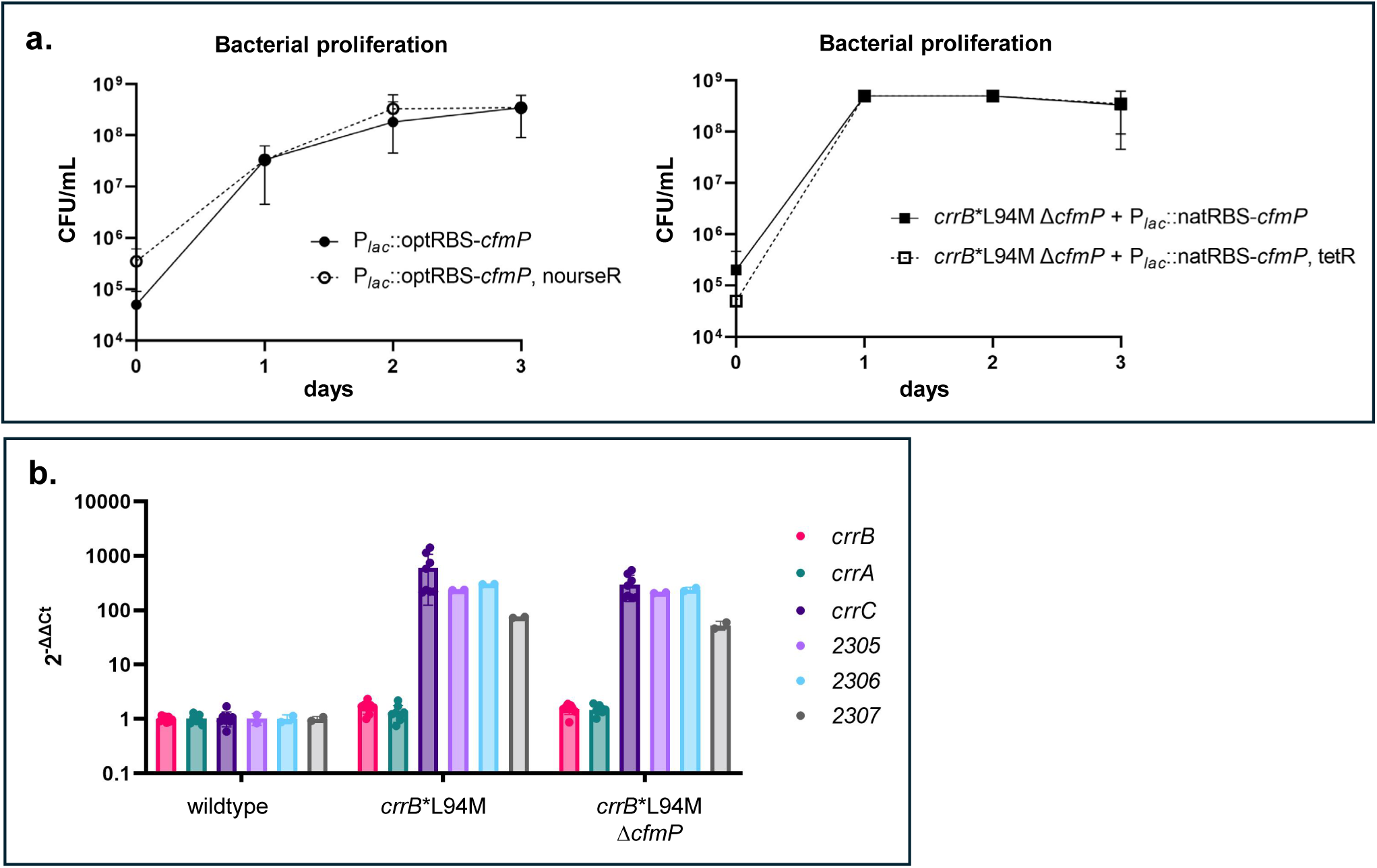
*cfmP* complementation and over-expression plasmids are maintained over the course of infection and Δ*cfmP* does not affect transcription of the *crr* locus. **(a)** Infected *G. mellonella* were plated on both KPC CHROMagar^TM^ to select for CR*Kp* and LB containing the antibiotic encoded on the *cfmP* overexpression (left, pUC19-P*_lac_*::optRBS-*cfmP*) or complementation (right, pACYC184-P*_lac_*::natRBS-*cfmP*) plasmids. Equal amounts of bacteria grew on both plates, indicating plasmids were maintained in the strains during infection. **(b)** RT-qPCR results of indicated strains show no changes in upregulation of the *crr* locus in *crrB*\*L94M versus *crrB*\*L94M Δ*cfmP,* indicating that changes in polymyxin resistance and virulence in *crrB*\*L94M Δ*cfmP* are not due to polar effects on the transcription of the *crr* locus.

**Extended Data Fig. 3:**
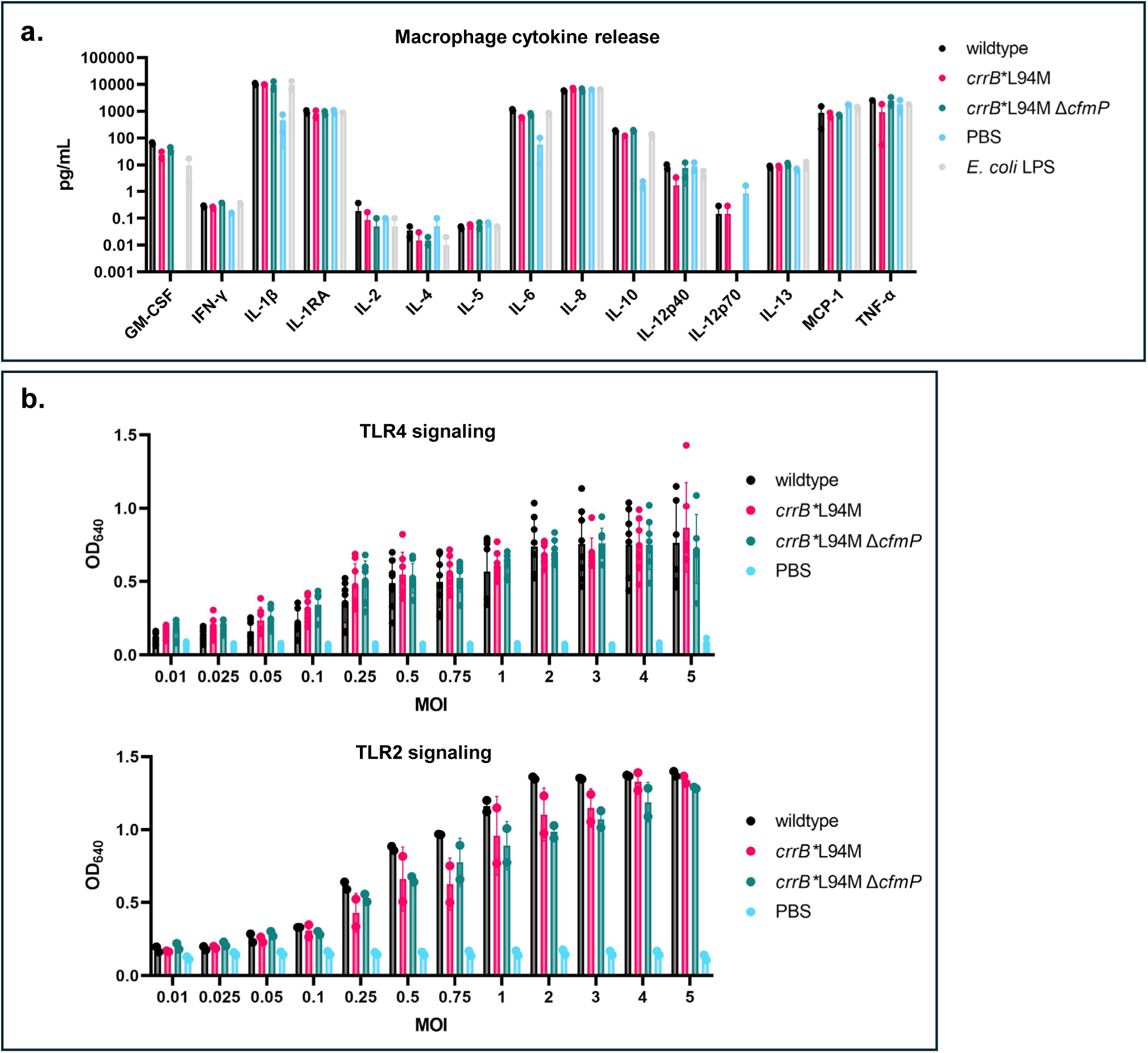
Innate immune signaling is unaffected by CrrB*L94M and CfmP. **(a)** Cytokine secretion in differentiated human macrophages in response to incubation with wildtype, *crrB*\*L94M, and *crrB*\*L94M Δ*cfmP* strains. Bacteria were grown to mid-log in CAMHB at 37°C and normalized to OD_600_=1.25 in 1X PBS. Bacteria were added to differentiated THP-1 monocytes at a multiplicity of infection (MOI) of 10 and purified LPS from *E. coli* 0111:B4 was added to a final concentration of 100 ng/mL, as a positive control. After 4 hours of incubation at 37°C, the supernatant was collected, filter sterilized, and sent for analysis using a commercially-available human cytokine array (Eve Technologies). **(b)** TLR4 and TLR2 signaling in response to incubation with heat killed bacterial cells from wildtype, *crrB*\*L94M, and *crrB*\*L94M Δ*cfmP* strains. Bacteria were grown to mid-log in CAMHB at 37°C and normalized to OD_600_=0.03125 in 1XPBS, corresponding to the concentration appropriate for the highest MOI. Cells were then heat killed at 80°C for 20 minutes. Bacteria were diluted in 1X PBS to reach appropriate MOIs in equivalent volumes. Cells were added to HEK-Blue hTLR2 or HEK-Blue hTLR4 (Invivogen) cells suspended in HEK-Detection medium and incubated for 18h at 37°C prior to OD_640_ readings.

**Extended Data Fig. 4:**
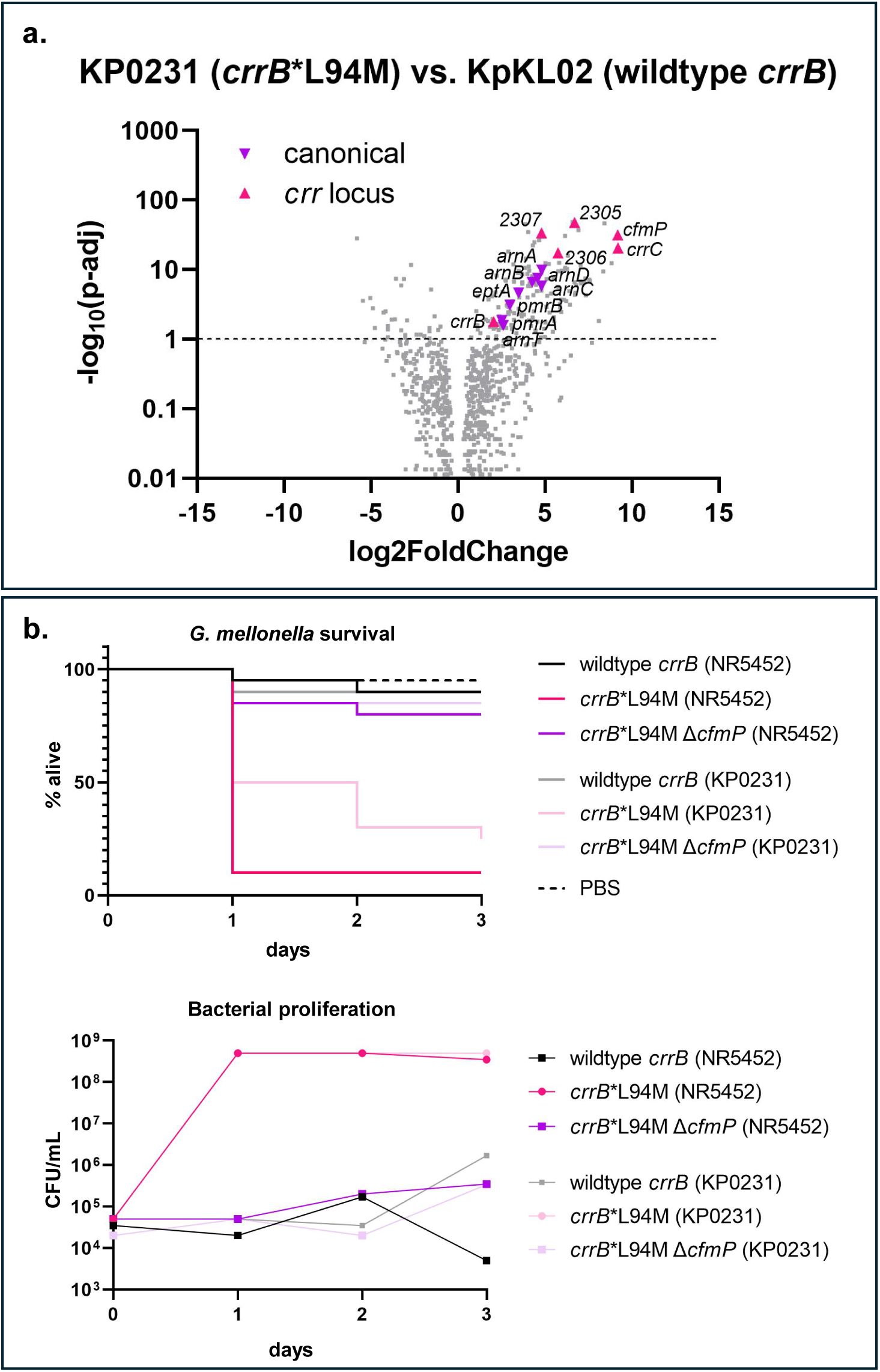
CrrAB-regulated genes and the virulence phenotype driven by CfmP are conserved in the original clinical isolate harboring *crrB*\*L94M. **(a)** Volcano plot of RNA-seq data generated from the original clinical isolate in which the *crrB*\*L94M mutation was identified (KP0231). The *crrB*\*L94M mutation was reverted back to wildtype *crrB* (KpKL02) and transcriptional profiles were compared. Genes in the *crr* locus and canonical genes involved in polymyxin resistance (*arn* operon and eptA) are highlighted. **(b)** Survival curve (top) and bacterial proliferation plot (bottom) of *G. mellonella* infections with the indicated two sets of isogenic strains. KP0231 is the original clinical isolate in which *crrB*\*L94M was identified. NR5452 is the parental wildtype *Kp* strain used for this study.

**Extended Data Fig. 5:**
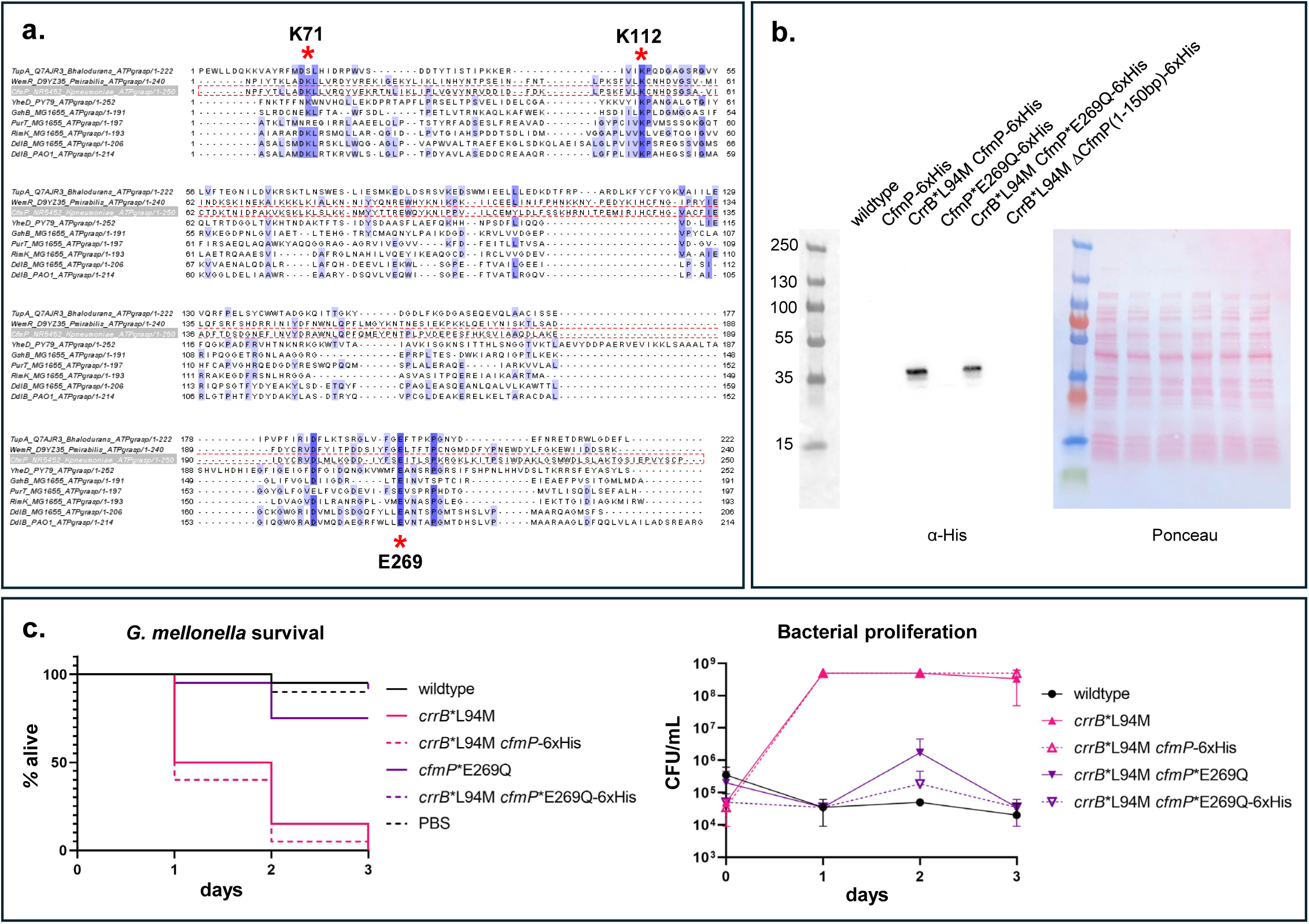
CfmP*E269Q is stable and 6xHis-tagged CfmP is functional. **(a)** Protein sequence alignment of the ATP-grasp domain of CfmP (amino acids 62-311) with the predicted ATP-grasp domain of WemR from *P. mirabilis*, and the bona-fide ATP-grasp domains of TupA (*B. halodurans*), YheD (*B. subtilis*), GshB (*E. coli*), PurT (*E. coli*), RimK (*E. coli*), and DdlB (*E. coli* and *P. aeruginosa*). Percentage identity is indicated in purple, and highly conserved residues involved in ATP binding are starred. **(b)** Immunoblot analysis of 6xHis-tagged protein levels in the indicated strains. Ponceau stain was used to control for loading. **(c)** Survival curves (left) and bacterial proliferation plots (right) of *G. mellonella* infections with the strains used for the immunoblot in (b).

**Extended Data Fig. 6:**
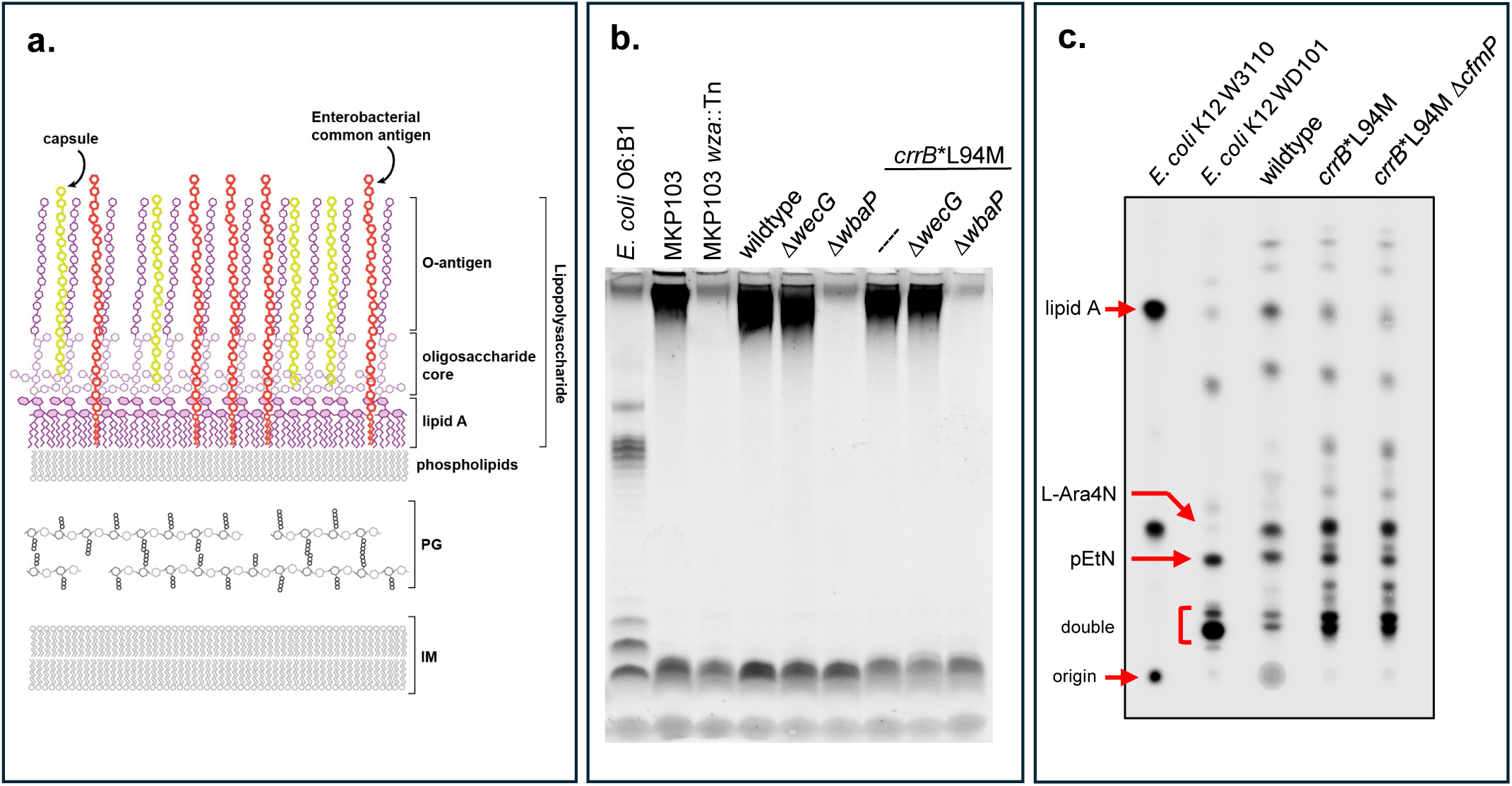
CfmP does not modify cell envelope polysaccharides or lipid A. **(a)** Schematic of the *Kp* cell envelope; PG: peptidoglycan layer, IM: inner membrane, outer membrane represented as the lipopolysaccharide and phospholipid asymmetric bilayer. **(b)** Emerald Pro-Q LPS stain of indicated strains. The wildtype *Kp* strain used in this study is lacking O-antigen, as demonstrated by the lack of O-antigen laddering. An *E. coli* O6:B1 strain was used as a control to indicate where O-antigen is detected on the gel. A deletion of *wbaP* (Δ*wbaP*) was introduced into wildtype *Kp* to prevent capsule synthesis and a deletion of *wecG* (Δ*wecG*) was introduced to prevent the synthesis of the Enterobacterial common antigen (ECA). The *Kp* strain MKP103 and its isogenic mutant with a transposon insertion disrupting the *wza* gene (MKP103 *wza*::Tn) necessary for capsule export, were used to indicate where capsule appears on the gel. **(c)** Thin layer chromatography (TLC) of purified lipid A isolated from the indicated strains. *crrB*\*L94M exhibits shifts consistent with the canonical aminoarabinose (L-Ara-4N) and phosphoethanolamine (pEtN) modifications added by the *arn* operon and *eptA* (both upregulated in *crrB*\*L94M). No change in the lipid A profile was observed when *cfmP* is deleted (*crrB*\*L94M Δ*cfmP*). *E. coli* K12 strains W3110 and WD101 were included as controls for unmodified and modified lipid A, respectively.

**Extended Data Fig. 7:**
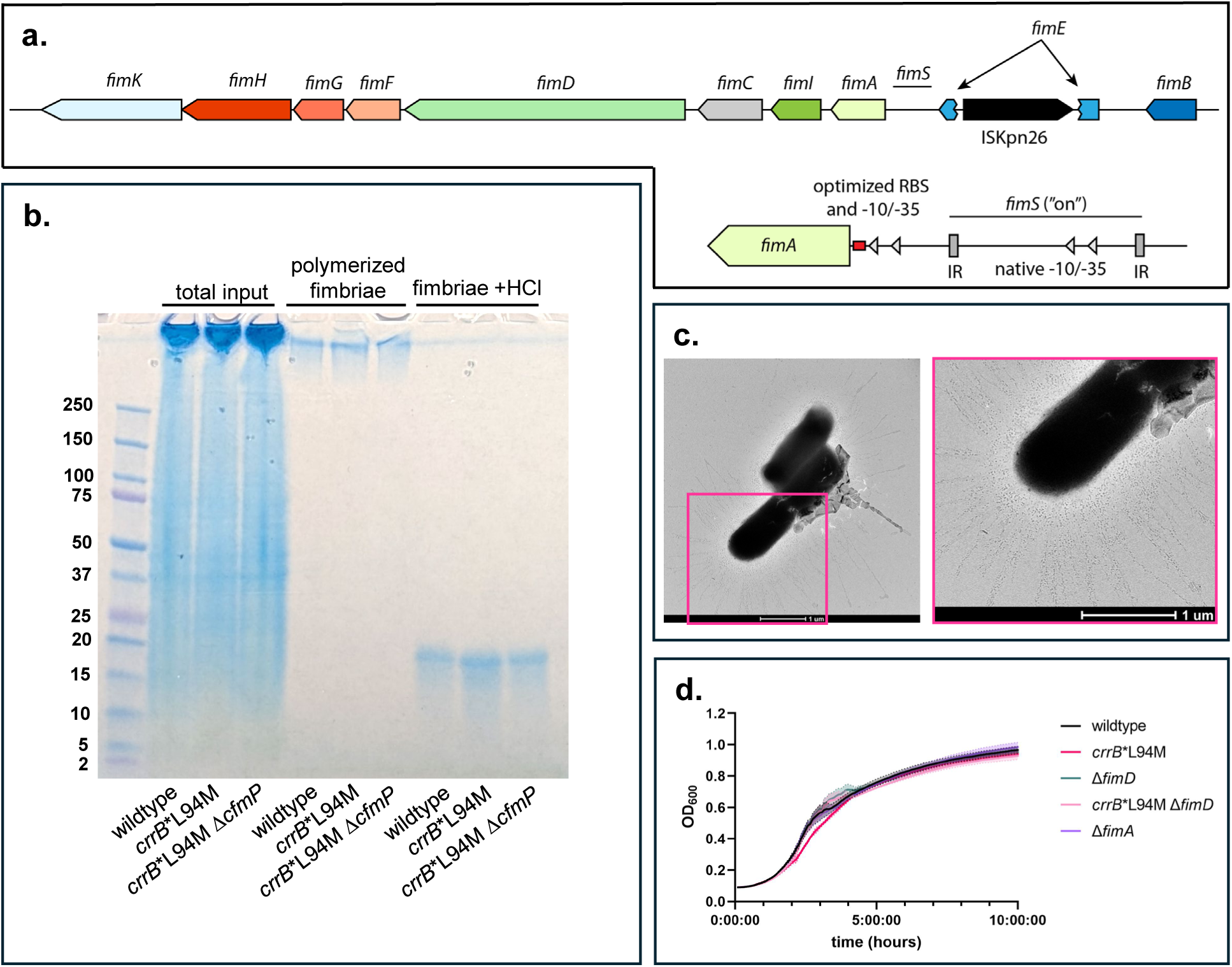
Fimbriae are produced during logarithmic growth in the wildtype *Kp* strain used in this study. **(a)** Schematic of the *fim* operon in the wildtype *Kp* strain used in this study. *fimE* coding region is interrupted by an insertion sequence (ISKpn26). Shown in the bottom right corner is the engineered promoter and ribosome binding site inserted upstream of *fimA* for constitutive expression of the *fim* operon. Also shown is the native *fimS* promoter region flanked by inverted repeats (IR). **(b)** Coomassie stained gel of fimbriae purified from wildtype, *crrB*\*L94M, and *crrB*\*L94M Δ*cfmP* strains. Polymerized fimbriae were treated with 1N HCl prior to boiling to depolymerize FimA and neutralized with 1N NaOH prior to loading on the gel. **(c)** Transmission electron micrograph of wildtype *Kp* grown to mid-log in CAMHB media; an enlarged inset is shown on the right, to show fimbriae polymers decorating the cell surface. **(d)** Growth curves of the indicated strains. Cells were grown to mid-log phase in CAMHB medium, normalized to an OD_600_ of 0.02 in CAMHB, and grown at 37°C. OD_600_ was measured every 5 minutes for 10 hours.

**Extended Data Fig. 8:**
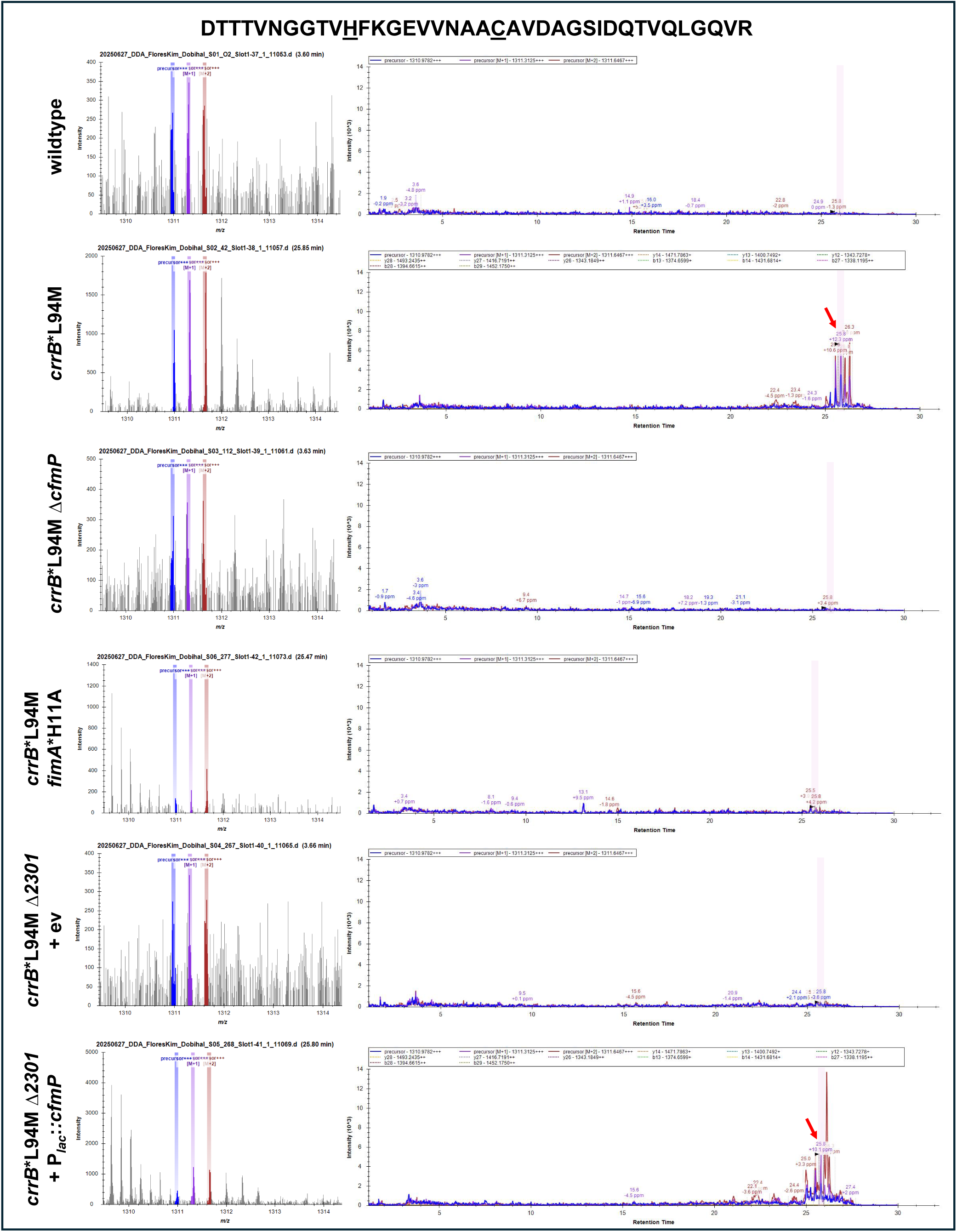
FimA peptide with oxidized H11 is CfmP-dependent. Peptide mass spectra (left) of the DTTTVNGGTVHFKGEVVNAACAVDAGSIDQTVQLGQVR peptide of FimA (left). LC-MS/MS chromatogram (intensity vs. retention time, right) of Skyline searches for that peptide containing oxidized H11 and the carbamidomethyl modification introduced during sample treatment (underlined in figure title). Proteomic data of purified fimbriae from each sample were initially processed in a non-specific search mode against a custom *K. pneumoniae* database containing FimA, FimF, FimG, and FimH sequences. The DTTTVNGGTVHFKGEVVNAACAVDAGSIDQTVQLGQVR peptide was identified in all samples. The same peptide with a +16 Da mass increase, consistent with oxidation of H11, was only identified in fimbriae samples isolated from *crrB*\*L94M and *crrB*\*L94M Δ*cfmP* containing the P*_lac_*::*cfmP* complementation vector (red arrows).

**Extended Data Fig. 9:**
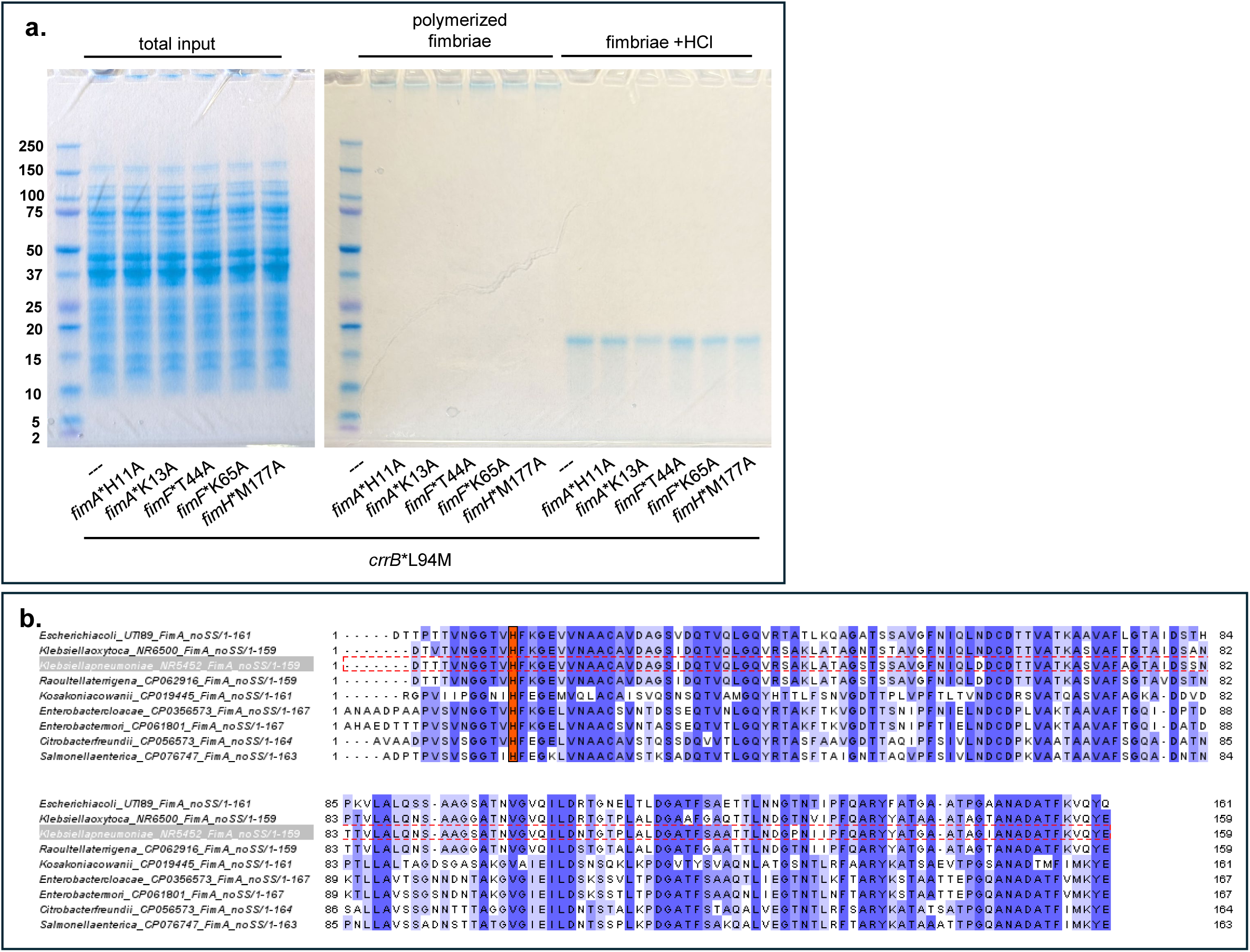
Fim* mutants are stably produced and FimA-H11 is conserved. **(a)** Coomassie stained gels of fimbriae purified from *crrB*\*L94M and *crrB*\*L94M *fim* mutants. Polymerized fimbriae were treated with 1N HCl prior to boiling to depolymerize FimA and neutralized with 1N NaOH prior to loading on the gel. **(b)** Multiple sequence alignments of FimA in *Kp* and other *Enterobacteriaceae* species containing *cfmP* and the *crr* locus. Genomes used for the alignment are the same as those found in Fig. 3b. The alignment is colored by sequence similarity, with the conserved histidine (H11) highlighted in orange. The sequence of *Kp* FimA is boxed in a dashed red line. FimA sequences were trimmed of their signal peptides prior to alignment.

**Extended Data Fig. 10:**
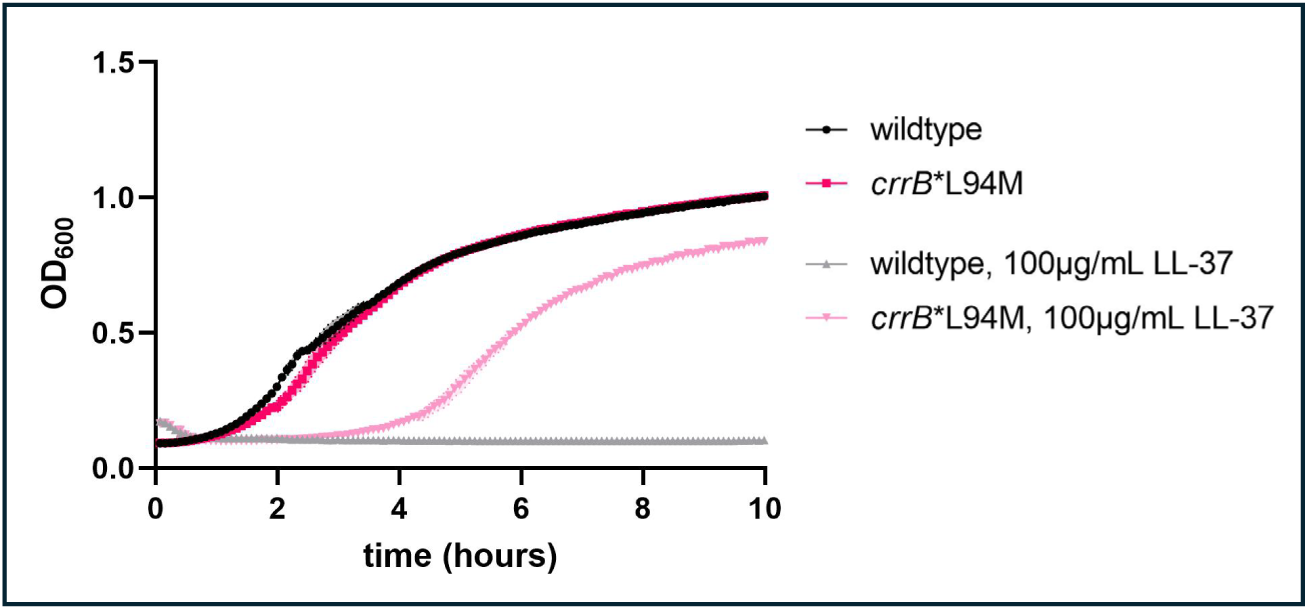
CrrB*L94M induces resistance to the human AMP LL-37. Growth curves of wildtype and *crrB*\*L94M *Kp*. Cells were grown to mid-log phase in CAMHB medium, normalized to an OD_600_ of 0.02 in CAMHB with and without 100 μg/mL LL-37, and grown at 37°C. OD_600_ was measured every 5 minutes for 10 hours.

